# Machine Learning Models Based on Histological Images from Healthy Donors Identify ImageQTLs and Predict Chronological Age

**DOI:** 10.1101/2025.09.04.674328

**Authors:** Ran Meng, William Zhu, Christopher JF Cameron, Pengyu Ni, Xiao Zhou, Tselmeg Ulammandakh, Mark B Gerstein

## Abstract

Histological images offer a wealth of data. Mining these data holds significant potential for enhancing disease diagnosis and prognosis, though challenges remain, especially in non-cancer contexts. In this study, we developed a statistical framework that links raw histological images and their derived features to the genotype, transcriptome, and chronological age of the samples. We first demonstrated an association between image features and genotypes, identifying 906 image quantitative trait loci (imageQTLs) significantly associated with image features. Next, we identified differentially expressed (DE) genes by stratifying samples into image-similar groups based on image features and performing DE comparisons between groups. Additionally, we developed a deep-learning model that accurately predicts gene expression in specific tissues from raw images and their features, highlighting gene sets associated with observed morphological changes. Finally, we constructed another deep-learning model to predict chronological age directly from raw images and their features, revealing relationships between age and tissue morphology, especially aspects derived from nucleus features. Both models are supported by a computational approach that greatly compresses gigapixel whole-slide images and extracts interpretable nucleus features, integrating both large-scale tissue morphology and smaller local structures. We have made all interpretable nucleus features, imageQTLs, DE genes, and deep-learning models available as online resources for further research.

**Significance Statement:** This study establishes a comprehensive framework that links histological image features to genotype, transcriptome, and chronological age in large-scale healthy tissue datasets, providing valuable insights into tissue morphology. By identifying 906 significant, interpretable imageQTLs and conducting differential expression analysis based on image features, we enhance understanding of genetic and morphological interactions. Additionally, we developed predictive models for both gene expression and chronological age from raw histological images, introducing a novel approach to studying age-related tissue-specific changes and presenting the first model to demonstrate the predictability of age from histological images.

## Introduction

Advanced histological image analysis, particularly when combined with neural networks and other analytical techniques, is an emerging field with the potential to improve both disease diagnosis and prognostic capabilities. Genotype, gene expression, aging, and environmental factors influence histological tissue morphology (*1–3*). Investigating the associations between histological image features and various levels of genetic data—such as genotype or transcriptomic data—across tissues from healthy donors is essential for uncovering the underlying molecular mechanisms that shape tissue morphology. Such insights are critical for understanding how disruptions in these mechanisms may lead to disease. Despite its importance, this area has not yet been fully explored. Here, we develop an approach that offers a hierarchical view of how genotype, gene expression, and epigenetic modifications shape histological morphology. As the global population ages, studying the relationship between histological images and aging will be crucial for explaining the mechanisms behind aging-related chronic conditions, such as cardiovascular disease, diabetes, Alzheimer’s disease, and certain cancers like prostate and lung cancer (*4*). Uncovering these basic processes can illuminate disease mechanisms, paving the way for better diagnosis and prevention.

However, multiple challenges remain. First, most studies have primarily focused on cancer or predicting gene expression in cancerous tissues, where histological images exhibit more distinctive features (*5*, *6*). For example, commonly observed features include a large nucleus with irregular size and shape, prominent nucleoli, scarce cytoplasm, and intense staining (*7*). These features limit the applicability of past models in non-cancerous contexts, which tend to have more subtle features. Another study applied a model in a non-cancerous setting, attempting to predict gene expression from histology images across multiple tissues. However, that study used a single, non–tissue-specific model. As a result, the predictions reflected average gene expression patterns across samples from each tissue, capturing general tissue-specific trends rather than precise, sample-specific expression profiles (*8*).

In addition to cancer or gene expression prediction, prior studies have attempted to address the association between genotype and histological image features, but they have either focused on a single tissue type (*1*) or used uninterpretable image features generated through dimensionality reduction techniques (*2*).

Moreover, interactions among genotype, gene expression, epigenetic modifications, age, and histological morphology are complex and multifaceted (**Fig. 1A**). Genetic material inherited from parents affects gene expression, which in turn impacts tissue morphology (*9*). The **aging** process plays a crucial role in **gene expression**, exhibiting tissue-specific effects (*10*). Yamamoto et al. (2022) showed that **aging** has a greater impact on gene expression patterns than **genetics** in most tissues, with age-related expression patterns being more tissue-specific (*10*). These researchers also demonstrated that most tissues align with Medawar’s hypothesis, where purifying selection (which eliminates harmful or deleterious mutations from a population) has a greater impact on early-life gene expression, except in highly proliferative tissues like blood and lung. Age has also been shown to be associated with **histological changes** in various tissues, including the brain and skin (*11*, *12*). In this paper, ‘age’ refers specifically to chronological age, which differs from ‘longevity,’ the total lifespan of an organism influenced by both genetic and non-genetic factors (*13*). In addition to these factors, **environmental stress** responses can lead to metabolic changes and epigenetic alterations, which may subsequently impact transcription. Environmental stress may also lead to somatic mutations, which can affect **gene expression, histological phenotypes**, disease susceptibility, and aging (*3*, *14*).

**Figure 1.**
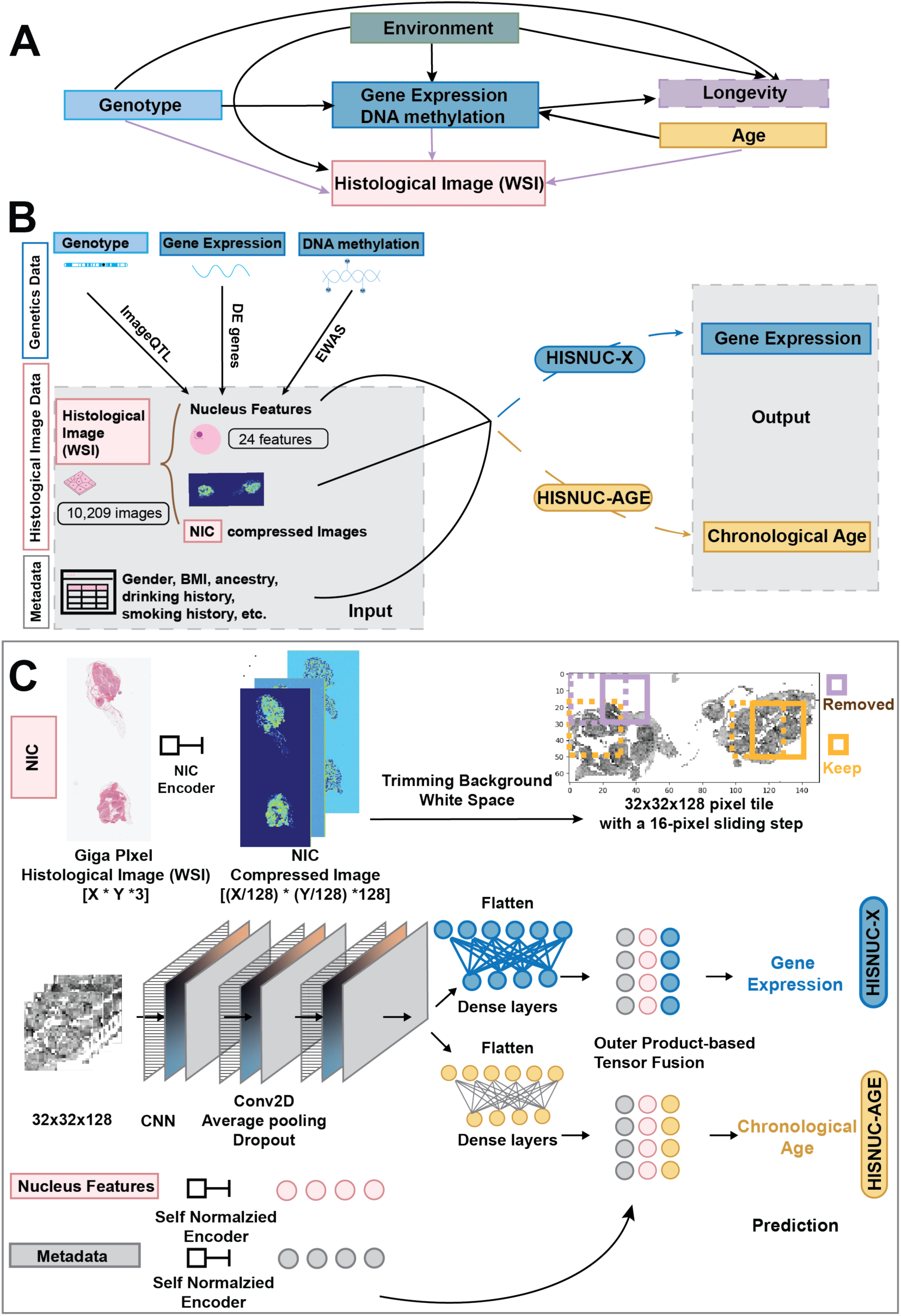
Dataset overview and network architecture. (A) Complex and multifaceted interactions among genotype, gene expression, histological images, age, longevity, and environment. (B) Data modality overview. This study focused on 12 tissues from the GTEx dataset, incorporating three key data types: genetic data, histological image data, and metadata. Nucleus features and NIC-compressed images were derived from the histological image data. Analyses such as imageQTL, DE analysis, and EWAS were conducted to identify associations between genetic data and interpretable image features. Nucleus features, compressed images, and metadata were used as inputs for both the HISNUC-X and HISNUC-AGE models. (C) Overall network architecture for the HISNUC-X and HISNUC-AGE models. The giga-pixel images (X*Y*3 pixels) were compressed into ((X/128) *(Y/128) *128) dimensions using the NIC algorithm. Background white space was trimmed, and the compressed WSI was tiled into multiple 32×32×128 tiles. Eight tiles with high tissue occupancy were then selected as input to HISNUC-X or HISNUC-AGE. Nucleus features and metadata were encoded using self-normalizing neural networks and then integrated with image features via tensor fusion to predict chronological age and gene expression (see **Methods** and **SI Appendix, Table S1**).

Previous studies have paved the way for utilizing deep learning to extract features for downstream data analysis (*15–17*). For example, the HE2RNA (*15*), Pacpaint (*16*), and PC-CHiP(*17*) models have demonstrated the ability to predict gene expression from histological tumor images or classify cancer subtypes. These studies have advanced the field of histological image analysis for predicting cancer-related content. However, features extracted by transfer learning, especially when pre-trained on non-histological datasets, may not transfer well to the specific and complex patterns seen in histology, leading to suboptimal performance and potential loss of critical information (*18*). While some models have been pre-trained specifically for histological image classification, they primarily focus on distinguishing tissue types through hierarchical feature extraction. Adapting these models for inter-individual predictions—such as predicting an individual’s gene expression in a specific tissue—may require specialized fine-tuning, as the pre-trained features may not fully capture the subtle variations necessary for accurate individual-level predictions (*18*).

In this study, we compressed gigapixel histological images to highly compact image representations using Neural Image Compression (NIC) (*19*). The NIC model is based on unsupervised image representation learning. Specifically, the NIC BiGAN (Bidirectional Generative Adversarial Network) encoder is pre-trained on histology images via adversarial learning, enabling the compression of gigapixel images into compact latent representations that capture deep semantic information. Instead of extracting a linear set of features, it reconstructs images, helping to preserve complex morphological details. It demonstrates exceptional performance and preserves the original spatial arrangements after compression. In addition to compressed images, we used QuPath (*20*) to extract interpretable image features of the nucleus, including nucleus area, eccentricity, circularity, and the nucleus-to-cell area ratio. We investigated the associations among these nucleus features, genotypes, and the transcriptome through Image Quantitative Trait Loci (imageQTL) and differential expression (DE) analysis. We identified 906 novel imageQTLs. These imageQTLs bridge the gap between genotype and image phenotype by illuminating the impact of genetic variations on interpretable image features. To further categorize the samples based on nucleus features, we identified DE genes by stratifying the samples according to nucleus characteristics (e.g., comparing samples with large vs. small nucleus area). Additionally, we examined the relationship between nuclear morphology and age by calculating the correlations between chronological age and nucleus features. Finally, by integrating compressed HIStological images, derived NUCleus features, and demographic meta-information, we developed two types of deep-learning models capable of accurately predicting tissue-specific gene eXpression profiles (HISNUC-X) and chronological Age (HISNUC-AGE) for healthy donors. With the HISNUC-AGE model, the histological morphology of skin, tibial nerve, tibial artery, and testis tissues exhibited greater predictive power for chronological age due to more pronounced age-related changes. Compared to previous histological image models that largely depend on uninterpretable image features, our model enhances explainability by using interpretable nucleus features and maintains spatial context within tissue morphology via compressed images.

## Results

### Dataset Overview and Overall Network Architecture

Our analysis leveraged data (and metadata) from 838 donors within the Genotype-Tissue Expression (GTEx) dataset (*21*) (**SI Appendix, Table S1** and **Fig. S1**), spanning 12 tissues and multiple data modalities, including genotype data, tissue-specific gene expression profiles, methylation sites, 10,209 histological images, and 24 distinct nucleus feature types (**Fig.1B**). Each GTEx tissue RNA expression sample was obtained through bulk sequencing and paired with a histological whole-slide image (WSI).

For each tissue type, we used QuPath to segment cells and nuclei and extract relevant nucleus features from WSIs (see **Methods**). These features included the nucleus area, circularity, eccentricity, and the nucleus-to-cell area ratio (**Fig. 2A**). Each WSI contained over 10,000 cells. We employed statistical measures—such as Q1, Q2, Q3, mode, mean, and standard deviation— to characterize the distribution of nucleus features (**SI Appendix, Dataset S1**). These nucleus features were integrated with genotype data from the donors, resulting in the identification of 906 imageQTLs, revealing associations between genetic variations and interpretable image features in particular tissues (**SI Appendix, Dataset S2** and **Dataset S3**). Additionally, we examined the associations between the transcriptome and nucleus features through DE analysis.

**Figure 2.**
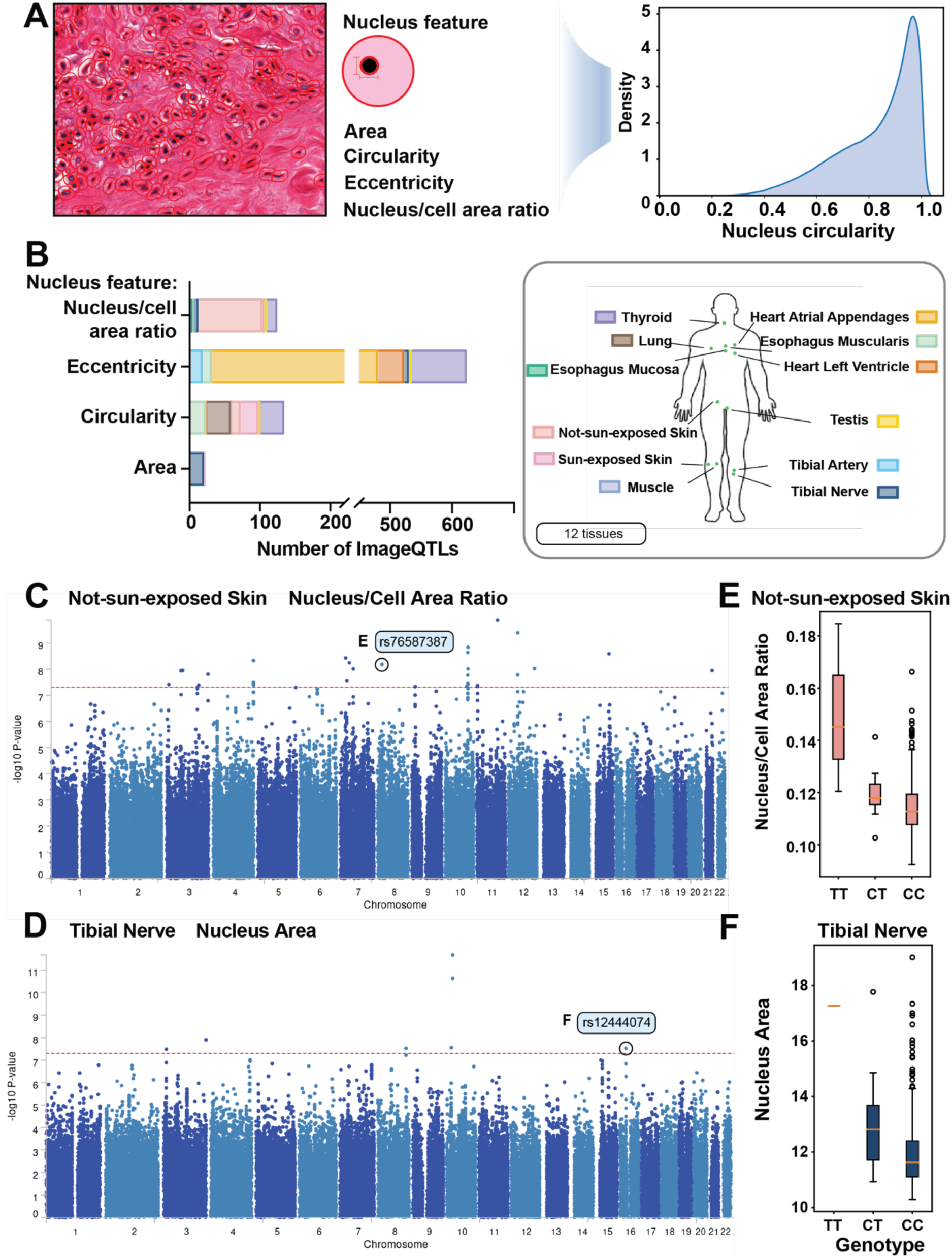
Associations among interpretable nucleus features, genotype, and gene expression. (A) An example image from the thyroid illustrates the image features identified by QuPath. (Left) The inner red circles represent nuclei segmented by QuPath, while the outer red circles delineate cell boundaries. This study focused primarily on nucleus area, nucleus circularity, nucleus eccentricity, and the nucleus-to-cell area ratio. (Right) Kernel density estimation of the distribution of nucleus circularity from the thyroid sample WSI. (B) Number of imageQTLs identified for each tissue and nucleus features. The box plot is color-coded according to 12 tissue types on the right. (The color scheme is consistent with that used in the other figures.) (C-D) Manhattan plots show imageQTL results for two representative tissues: (C) not-sun-exposed skin tissue with the nucleus-to-cell area ratio and (D) tibial nerve tissue with the nucleus area. The Y-axis represents -log_10_ P-values, while the X-axis displays SNP loci across chromosomes. Circled dots highlight two SNPs shown in panels E and F. (E) Box plot illustrating the nucleus-to-cell area ratio across the reference and alternative alleles of rs76587387 in the not-sun-exposed skin tissue. (F) Box plot depicting the distribution of nucleus area across the reference and alternative alleles of rs12444074 in tibial nerve tissue.

For each tissue, we trained two convolutional neural network (CNN)-based multimodal models to predict: (1) gene expression (HISNUC-X) and (2) chronological age (HISNUC-AGE) (**Fig. 1C**). Both models employ a CNN architecture to extract features from compressed WSIs. Nucleus features (extracted via QuPath) and donor metadata were encoded using self-normalizing neural networks (SNNs)(*22*), and all modalities were integrated through a fusion layer (*23–25*)(see **Methods** and **SI Appendix, Table S1** for details). We have made the extracted nucleus features, imageQTLs, DE genes, and both sets of pretrained models publicly available as data resources (see **SI Appendix and Data, Materials, and Software Availability**).

### Analyzing Image Feature Associations with Genotype to Derive ImageQTLs

By integrating nucleus features extracted via QuPath with GTEx genotype data, we identified associations between single-nucleotide polymorphisms (SNPs) and individual image features, uncovering 906 SNP loci significantly associated with image traits (imageQTLs) (**SI Appendix, Dataset S2)**. After linkage disequilibrium pruning and functional annotation using FUMA, these loci were consolidated into 384 lead SNPs (see **Methods** and **SI Appendix, Dataset S3**).

The image features most frequently associated with imageQTLs were nucleus eccentricity and circularity. The number of imageQTLs per tissue is shown in **Fig. 2B**. Notably, the skin (not-sun-exposed), heart atrial appendage, and thyroid tissues exhibited relatively large numbers of imageQTLs (**Fig. 2B**). Although the sun-exposed skin tissue had a larger sample size than its not-sun-exposed counterpart (**SI Appendix, Table S1**), which typically provides greater power in QTL calculations, the number of significant SNPs associated with the sun-exposed skin did not exceed that of the not-sun-exposed tissue. This result is likely due to the greater environmental impact on the sun-exposed tissue.

Among the imageQTL loci, 594 genes were mapped using FUMA’s SNP-to-gene annotation (**SI Appendix, Dataset S3)**. Of these, we identified a list of genes with potential roles in regulating nuclear morphology, based on experimental evidence or literature-based inferences suggesting plausible mechanistic links (**SI Appendix, Dataset S4**). Here, we present strong examples of imageQTL SNPs linked to morphological changes through well-established biological mechanisms. First, as shown in **Figs. 2C** and **2E**, SNP rs76587387 (on chromosome 8—chr8) was correlated with the nucleus-to-cell area ratio in not-sun-exposed skin tissue. The alternative allele was associated with a larger nucleus-to-cell area ratio. This SNP maps to the *LZTS1* gene (**SI Appendix, Fig. S2)**, which encodes a nuclear-localized tumor suppressor involved in microtubule stabilization, mitotic progression, and cell growth regulation, and also influences the endosome–nucleus distance (*26*). Given the role of mitotic control in regulating nucleus size, we propose that allelic variation at this locus may alter nuclear morphology via modulation of *LZTS1* mediated cell cycle progression or endosome–nucleus positioning (**SI Appendix, Fig. S3**). Second, SNP rs12444074 (on chr16) (**Figs. 2D** and **2F)** was correlated with the “nucleus size” feature in tibial nerve tissue, showing significant differences between the alternative and reference alleles. The reference allele was associated with a smaller nucleus size. This SNP maps to *KIAA0556* (**SI Appendix, Fig. S2)**, which encodes a microtubule-associated ciliary base protein and is also localized close to the nucleus (*27*). Its known functions—anchoring microtubules and coordinating nucleus–cytoskeleton interactions—suggest a role in modulating nucleus size and positioning (**SI Appendix, Fig. S3**). These findings demonstrate that imageQTLs capture associations between genotype and interpretable nucleus features in histological images.

In addition, we identified 50 imageQTLs that overlap with GTEx expression quantitative trait loci (eQTLs) (see **Methods** and **SI Appendix, Dataset S5**), suggesting that these variants may influence nucleus features by modulating the expression of associated eGenes. One such example is ***GSDMD***, a gene implicated in both imageQTL and eQTL analyses. This gene (encoding gasdermin D), activated by inflammatory caspases, is a key effector of pyroptosis. Upon activation, its N-terminal fragment (GSDMD-NT) forms membrane pores—primarily at the plasma membrane, but recent studies suggest it can also target intracellular membranes, including the nuclear envelope (**SI Appendix, Fig. S3**). This disruption of membrane integrity may permit nuclear entry of cytosolic factors such as caspase-11, which has been implicated in histone cleavage and chromatin remodeling, and ultimately may alter nuclear morphology observable by microscopy (*28*, *29*).

### Linking Nuclear Morphology to Differential Gene Expression and DNA Methylation

To investigate the association between image-derived nucleus features and gene expression, we stratified samples based on their nuclear characteristics and identified DE genes between groups (e.g., large vs. small nucleus area or high vs. low nucleus-to-cell area ratio) (**Fig. 3**; see **Methods** and **Data, Materials, and Software Availability**). In particular, we found that *LMNA* and *ACTB* were differentially expressed in association with several nucleus features across multiple tissues. High expression of *LMNA* and *ACTB* was associated with an increase in nucleus size. Additionally, high *LMNA* expression was evident in clusters with greater nucleus eccentricity. Previous studies showed that *LMNA* encodes lamin A/C proteins, which are associated with nucleus size, and that overexpression of *LMNA* leads to an increased nucleus aspect ratio (*30*). The *ACTB* gene encodes the beta-actin protein, which plays a role in the structure and shape of the cell and also affects nucleus shape (*31*).

**Figure 3.**
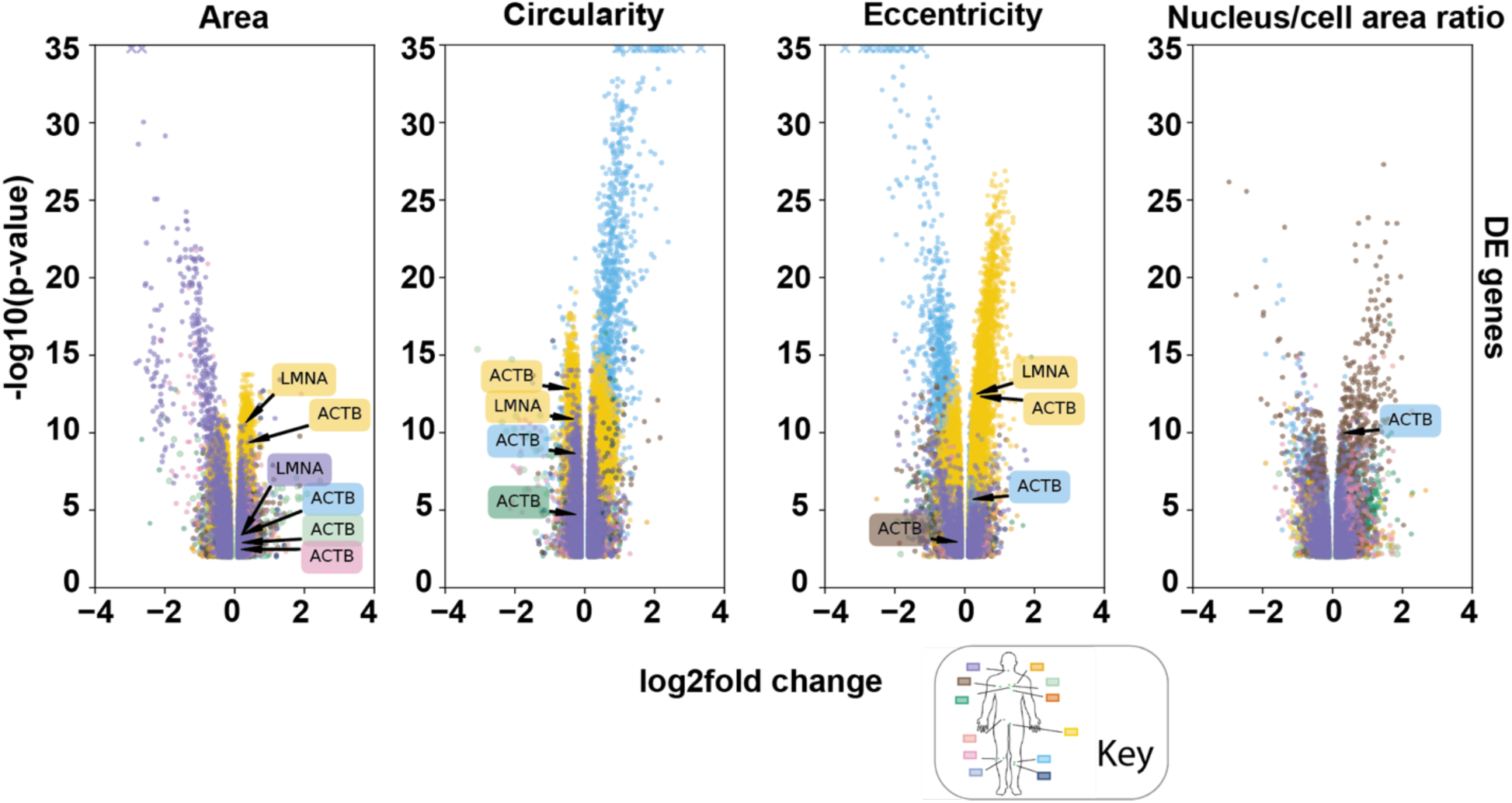
DE analysis for different nucleus feature groups. Log_2_ fold changes and adjusted p-values for DE genes based on various nucleus features. Each dot represents a gene. Dots are color-coded according to tissue types. Genes with a -log_10_(adj. p-value) greater than 35 are marked with an ‘x’. Genes with an absolute log_2_ fold change greater than 0.1 and an adjusted p-value less than 0.01 are shown in the figure.

Additionally, we compared genes mapped by imageQTL SNPs to the DE genes, revealing a subset of overlapping genes (**SI Appendix, Dataset S6**). This overlap suggested that certain SNPs and their adjacent genes may directly influence nucleus features or co-regulate these features through alterations in gene expression. Three good examples of this are *TMPRSS2*, *CD2AP*, and *FRY*(*32–35*). These genes have supporting evidence from previous studies demonstrating that changes in their expression or function can lead to alterations in nucleus shape, size, or structure (**SI Appendix, Dataset S7**).

In the above sections, we explored the relationship between nucleus features, genetic variants (SNPs), and gene expression. Next, we investigated whether nucleus features are also associated with DNA methylation, which is generally considered an upstream regulator of gene expression. To do this, we conducted an epigenome-wide association study (EWAS) focused on lung tissue data from eGTEx (*36*, *37*). The analysis focused solely on lung tissue, as this represents the largest methylation dataset available in the eGTEx database. We correlated nucleus features with methylation values at each CpG site, identifying methylation sites significantly associated with nucleus features such as nucleus eccentricity and the nucleus-to-cell area ratio (see **Methods**, **SI Appendix, Fig. S4,** *left,* and **Data, Materials, and Software Availability**). Gene set enrichment analysis (GSEA) using these sites revealed substantial overlap between the enriched gene sets and the DE genes related to lung nucleus features (**SI Appendix, Fig. S4,** *right*).

### HISNUC-X Model: Tissue-Specific Prediction of Gene Expression

To further explore the relationship between gene expression and histological images, we developed 12 tissue-specific models to predict gene expression based on WSIs. We defined tissue-specific genes as those with a Tau score greater than 0.85 (*38*) and gene expression levels greater than 1. The number of tissue-specific genes identified for each tissue varied, ranging from 156 to 6,677 (**Fig. 4A**). The testis had over 6,500 tissue-specific genes, considerably more than other tissues. For each tissue model, the 100 genes with the highest expression levels were selected as targets. To link nuclear morphology to gene expression, we calculated the Pearson correlations between nucleus features and the 100 selected genes. For instance, **Fig. 4B** depicts the correlation between nucleus features and 100 tissue-specific gene expression profiles in lung tissue. We evaluated the relationship between nucleus features and gene expression and subsequently used these features as input to the HISNUC-X model (**SI Appendix, Dataset S8,** details in **Methods**).

**Figure 4.**
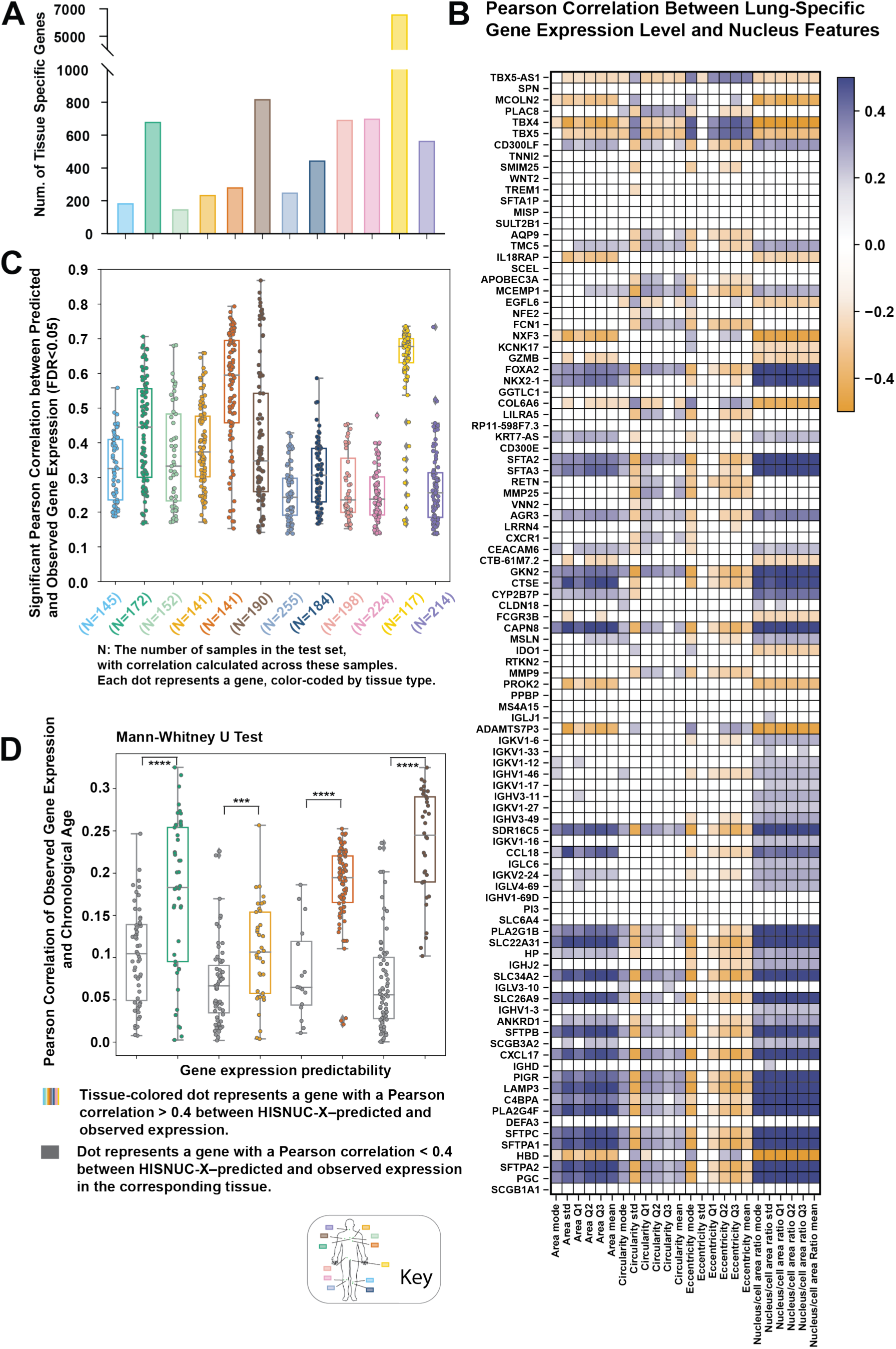
Prediction of tissue-specific gene expression using HISNUC-X. (A) The number of tissue-specific genes identified using the Tau score. (B) Pearson correlation between nucleus features extracted by QuPath and tissue-specific gene expression levels in the GTEx lung dataset. (C) Pearson correlation (adj.p <0.05) between predicted and target gene expression levels in the test set. The distribution of correlations for all predictable genes is presented in box plots. (D) Pearson correlation between observed gene expression level and chronological age across samples. Each dot represents one gene, with the Y-axis displaying the correlation. For each tissue, genes were categorized into two groups based on whether the Pearson correlation between HISNUC-X–predicted and observed expression exceeded 0.4. Genes with correlation > 0.4 are color-coded by tissue type, while those with correlation ≤ 0.4 are shown in grey. We conducted the Mann–Whitney U test (one-sided) to compare the distributions of correlations between these two groups. The results demonstrated a significant difference in the age-correlation distributions between high- and low-prediction-accuracy genes.

For each prediction target, we assessed the correlation between predicted and observed gene expression levels using the held-out test set. **Fig. 4C** highlights the Pearson correlation for each gene in each tissue, focusing on predictable genes—those with significant correlations across three independent experiments. Significance was determined using Fisher’s method (*39*) followed by the Holm–Šidák (HS) method to control for multiple testing (adjusted p-value < 0.05; **SI Appendix, Dataset S9**). Median absolute error (MAE) is also reported for each gene prediction. Notably, genes from skin tissues were less predictable compared to those from esophagus mucosa, heart, lung, tibial nerve, and testis tissues. Within the same tissue, certain genes demonstrated predictable expression, while others did not. To gain deeper insights into this variability, we investigated why some tissue-specific genes were more predictable than others in the same tissue. In the four somatic tissues with the highest gene-level predictability—esophagus mucosa, heart atrial appendage, heart left ventricle, and lung—we observed both strong prediction correlations and a large number of predictable genes. Among these four tissues, genes with a Pearson correlation > 0.4 between HISNUC-X–predicted and observed expression generally showed significantly stronger associations with chronological age compared to those with correlations ≤ 0.4 (**Fig. 4D**). To test whether gene expression predictability reflects an underlying association with age—even when age is not used as an input—we retrained the model shown in **Fig. 4D** without including the ‘age’ variable in the metadata.

### HISNUC-AGE Model: Tissue-Specific Prediction of Chronological Age

We trained HISNUC-AGE models for 12 different tissues to predict chronological age, integrating compressed WSIs, interpretable nucleus features, and donor metadata as inputs. All models followed the network architecture in **Fig. 1C**, with slight variations in WSI tiling size and learning rate (details in **SI Appendix, Dataset S10**).

The nucleus features were evaluated for their correlation with age (**Fig. 5A**). In particular, the features in lung, tibial nerve, sun-exposed skin, and testis were especially strongly correlated. Nucleus-to-cell area ratios showed a negative correlation with chronological age across most tissues. In contrast, the variability of nucleus circularity increased with age in most tissues, indicating greater heterogeneity in nucleus shape in older age (*40*), and nucleus eccentricity (Q1, Q2, Q3, mean, mode) showed a positive correlation with age in most tissues. The latter finding is supported by Iijima et al. (2023), who reported that increased nucleus eccentricity with age is associated with decreased α-Klotho expression (*41*).

**Figure 5.**
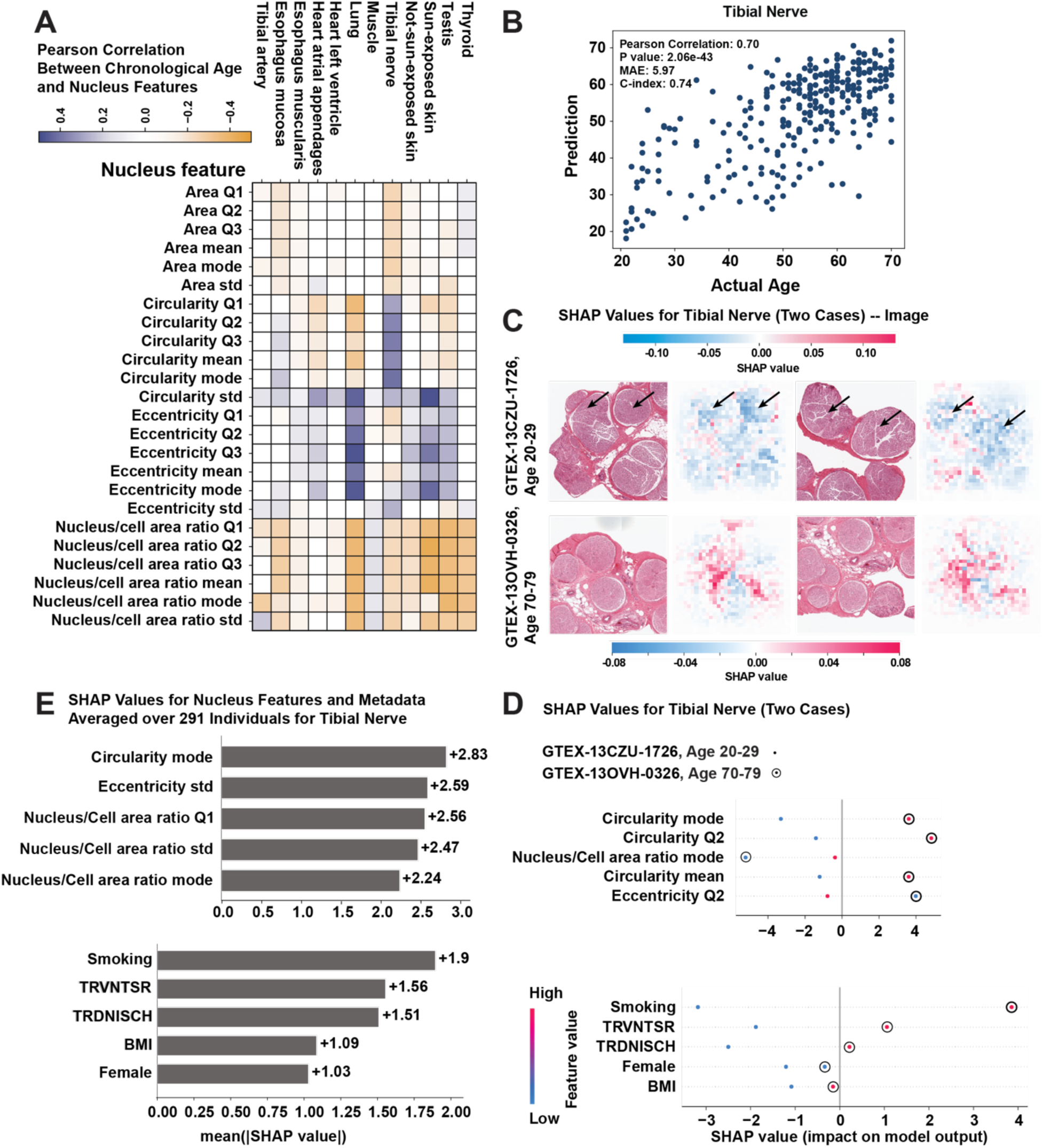
Chronological age prediction for tibial nerve using the HISNUC-AGE model. (A) Pearson correlation between chronological age and QuPath-extracted nucleus features for each GTEx tissue dataset. (B) The scatter plot shows that the predicted age and the actual chronological age for tibial nerve samples from a single prediction experiment. The correlation is 0.70 (p-value 2.06e-43). The MAE is 5.97 years between predicted and target values. (C) SHAP analysis highlights the pixel weights mapped back to the original image for two samples from different age groups, with positive SHAP pixels shown in red (contributing to older age predictions) and negative SHAP pixels shown in blue (contributing to younger age predictions). Arrows indicate the fiber regions. Fig. 5C (*top*) presents two tiles from the WSI of the 20–29-year-old donor, while Fig. 5C (*bottom*) shows two tiles from the 70–79-year-old donor. In the younger donor’s tiles, myelinated fiber regions (highlighted in blue and indicated by black arrows) contribute to the younger age prediction, whereas in the older donor’s tiles, regions associated with older age (shown in red) are more prominent. (D) Comparison of SHAP values for nucleus features and metadata between two samples from different age groups. Dots are colored by feature value, from red (high) to blue (low). The SHAP values on the X-axis indicate their impact on the model output. Only the top 5 features (ranked by feature importance based on these two samples) that contribute to the model are displayed, distinguishing the two samples from different age groups. TRVNTSR indicates whether the donor was ventilated for less than 24 hours before the estimated procurement start time. TRDNISCH represents the total ischemic time for the donor. (E) Mean absolute SHAP values for vectorized features across the entire test set, ranked by feature importance.

**Fig. 5B** shows the chronological age prediction results for the tibial nerve. We used SHapley Additive exPlanations (SHAP) values to assess the feature importance of compressed images (**Fig. 5C**) (*42*), nucleus features, and meta-information for age prediction (**Figs. 5D and 5E**). For illustration, we present two representative examples from different age groups: a 20–29-year-old donor (GTEX-13CZU-1726) and a 70–79-year-old donor (GTEX-13OVH-0326) (**Figs. 5C and 5D**). The SHAP values highlight how the myelinated fiber regions in the younger donor are important for the age prediction. Moreover, this finding is supported by previous studies (*43*) showing that thicker myelin and larger fiber diameters are inversely correlated with aging in mice.

Furthermore, **Fig. 5E** explores the contributions of the top 5 nucleus features and top 5 meta features across the entire test set for the tibial nerve. Overall, nucleus features—such as the mode of nucleus circularity, the standard deviation of eccentricity, and the nucleus-to-cell area ratio—were the most significant contributors to chronological age predictions. Using a similar strategy, we also analyzed image, nucleus, and meta-feature contributions for sun-exposed skin, providing examples that demonstrate the model’s interpretability (**SI Appendix, Fig. S5**).

Stepping back to overall performance, **Fig. 6** displays the correlation between predicted and the actual chronological age from the test sets across all 12 tissues (**SI Appendix, Fig. S6; Datasets S10 and S11**). The MAE and concordance-index (c-index) were used to evaluate prediction accuracy (**SI Appendix, Dataset S11**). Four tissues demonstrated particularly strong predictive performance: tibial artery, tibial nerve, sun-exposed skin, and testis. However, the chronological ages of the two heart tissues were less predictable, likely because the biological age of the heart tends to be consistently younger than its chronological age (*44*).

**Figure 6.**
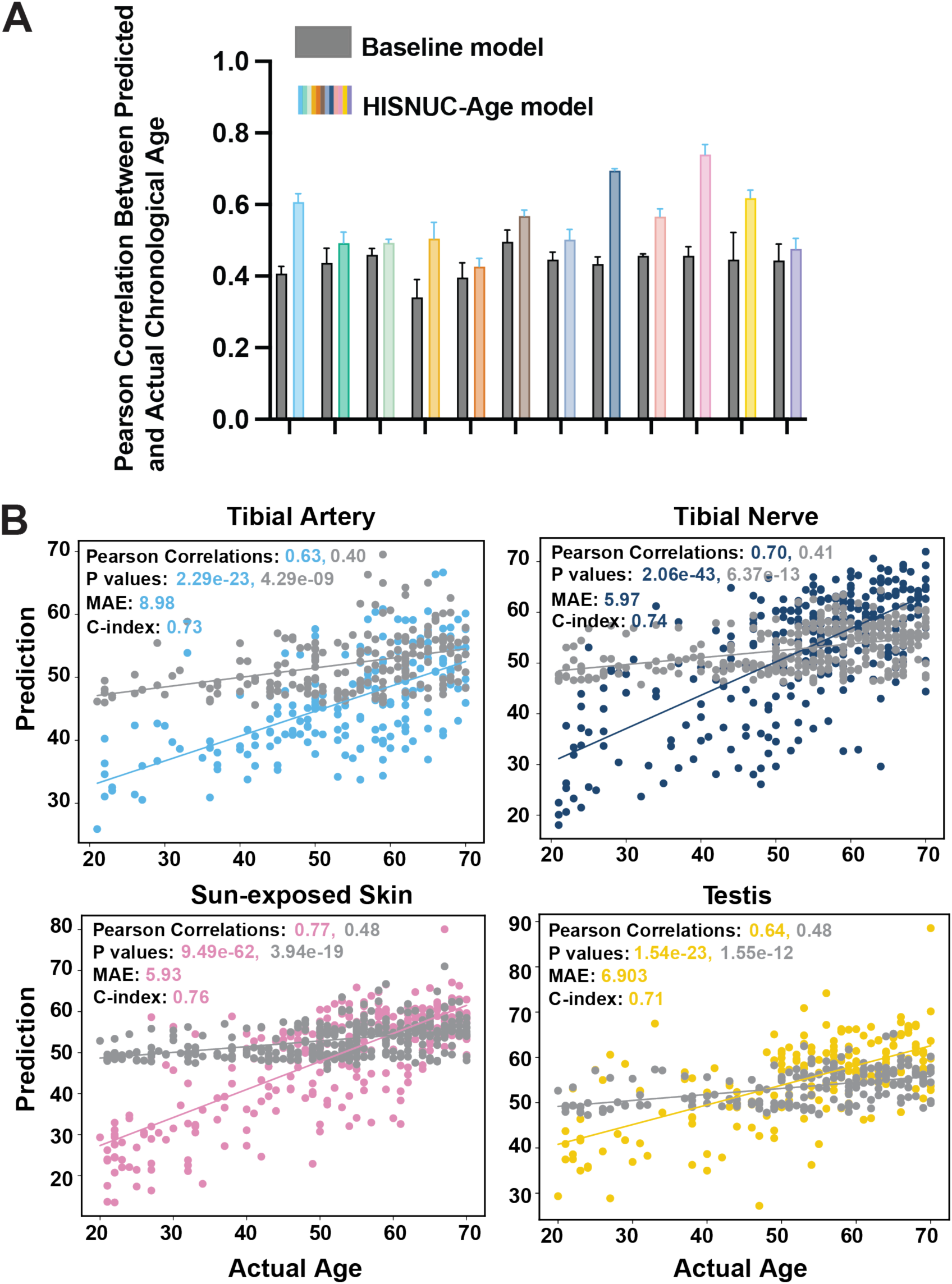
Correlation between chronological age and tissue-specific age predicted by the HISNUC-AGE model. (A) Comparison of correlation results across tissues shows elastic net baseline model predictions in grey and HISNUC-AGE results colored according to each tissue. The error bars represent one standard deviation above and below the mean, illustrating the spread of the data around the average from three independent prediction experiment replicates (see **SI Appendix, Datasets S11 and S13** for details). (B) Scatter plot of the predicted age vs. the actual chronological age for each tissue from a single prediction experiment, including the best-fit line. The scatter plot colored in grey depicts predictions from metadata using the elastic net baseline model as a control. The scatter plot colored by tissue displays predictions from the HISNUC-AGE models.

As a control, we employed an elastic net baseline model using only metadata as input (**SI Appendix, Datasets S12 and S13;** details in **Methods)**. Compared with the elastic net baseline model, the additional information from histological imaging improved aging predictions across all tissues.

## Discussion

Computational histopathology workflows play a crucial role in analyzing large gigapixel WSIs. We leveraged the NIC and QuPath algorithms to integrate interpretable nucleus features with histological images in a deep-learning context. Our HISNUC models effectively preserve large-scale tissue morphology and spatial information while capturing fine local nuclear structures. We applied QuPath to analyze 10,209 histology WSIs from various tissues in the GTEx dataset, detecting tens of thousands of nuclei per WSI and deriving their quantitative features. As the most efficient software available for large-scale histological imaging analysis, QuPath facilitated cell detection, segmentation, and feature extraction, producing valuable quantitative data on cell and nuclear morphology as well as staining intensities. These extracted image features enhanced interpretability in imageQTL analysis, DE analysis, and HISNUC models. Unlike previous studies that relied on abstract features to identify imageQTLs, our approach directly indicates which aspects are potentially affected by SNPs and genes. Incorporating interpretable image features across multiple tissues enhances our understanding of the intricate relationships among genetics, epigenetics, and tissue morphology. Additionally, the nucleus features detected for each image in the dataset, along with imageQTLs and DE genes, serve as valuable resources for future analysis.

Histological images at different scales emphasize distinct levels of morphological detail(*45–47*). In our study, we used GTEx WSIs scanned at 20× magnification (0.4942 μm/pixel) without downsampling, providing the highest available resolution. At this scale, a typical nucleus spans approximately 10–20 pixels in diameter, and small regions (20–60 pixels) can capture subcellular or cellular structures, depending on tissue type. Larger tissue-level features—such as glandular organization, stromal composition, and vascular structures—emerge across broader regions spanning thousands of pixels (e.g., 4,096×4,096) (*47*). In our models, QuPath-extracted nucleus features from 20× WSIs provide quantitative subcellular morphology, and ablation experiments confirm their critical importance—removing nucleus features significantly reduces model performance. Furthermore, the NIC-giga framework compresses raw 128×128 RGB patches into compact 128-dimensional vectors using an autoencoder pre-trained specifically on histopathology data to preserve high-level subcellular and cellular features. We then aggregate multiple 64×64 (or 32×32) compressed patches per WSI, effectively spanning raw image areas of up to 8,192×8,192 (or 4096×4096) pixels. By integrating multiple patches across each slide, the model learns diverse spatial contexts and captures tissue heterogeneity. This architecture enables the model to represent a broad morphological spectrum—from nucleus shape and size to tissue-level architecture and spatial organization. Ultimately, our multimodal approach combines fine-grained image features, cell-level morphological measurements, and patient-level metadata, capturing both micro- and macro-structural cues relevant to biological aging and transcriptomic variation. These inputs are fused through a deep neural network that enables robust prediction while maintaining interpretability through nucleus-derived features.

We recognize potential areas for enhancing our study. First, technical variations in histological image capture and processing may impact the detected features; for example, sample thickness and the slicing process may affect tissue shape and morphology. Comparing WSIs paired with bulk sequencing data to those paired with spatial transcriptomic data is akin to the distinction between bulk RNA sequencing and single-cell sequencing. While bulk methods are more cost-effective, they generally provide lower resolution. The nucleus image features extracted from each WSI may originate from various cell types. To mitigate these challenges, we employed statistical distributions to describe nucleus features within a WSI, converting raw feature data into structured inputs for integration with imaging and metadata in predictive modeling. In our imageQTL studies, we used the mode as a representative value to characterize the predominant image features within each WSI, illustrating how alternative alleles influence these nucleus features. Nevertheless, determining a more accurate and representative value for QuPath detection within each WSI remains an open question. By leveraging spatial transcriptomic data for training, we can partially address these issues, resulting in models that not only enhance prediction accuracy but also reduce the costs associated with spatial transcriptomics. For instance, a well-trained model can predict gene expression without the need for sequencing. This approach would significantly enhance the interpretation of image features extracted, establishing a vital connection with localized expression data.

## Methods

### Nucleus Feature Extraction

QuPath v0.4.3 was used to extract nucleus features from GTEx WSIs. QuPath is publicly available (https://github.com/qupath/qupath/releases). Default settings were followed as outlined in the QuPath tutorial: https://qupath.readthedocs.io/en/stable/docs/tutorials/index.html. The image type was set to ‘brightfield H&E,’ and stain vectors were optimized. Subsequently, we ran a pixel classifier to identify relevant tissue areas. Cell and nucleus detection was performed using QuPath’s cell detection tool. Finally, the extracted cell and nucleus detection measurements for each WSI resulted in a structured dataset for further analysis. We configured the pipeline for each tissue type to accommodate different tissue morphologies. This automated feature extraction was crucial for ensuring consistent and objective quantification of tissue characteristics across all slides.

Four key nucleus features were used in this study: (1) nucleus area, (2) nucleus circularity, (3) nucleus eccentricity, and (4) nucleus-to-cell area ratio. We excluded certain features, such as the nucleus perimeter and max caliper, due to their high correlation with nucleus area. Each WSI contained over 10,000 cells; therefore, we employed several parameters to characterize the distribution within each WSI. Outliers were filtered using a 1.5 interquartile range. We then applied kernel density estimation, partitioning the data into 500 bins, and selected the value corresponding to the highest density, representing the mode of the entire distribution. Additionally, we included the first quartile (Q1), second quartile (Q2), third quartile (Q3), mean (here we used a 90% trimmed mean), and standard deviation to describe the distribution. These six representative values for each nucleus feature were then utilized in subsequent analyses, resulting in 24 nucleus features in total.

### ImageQTL Analysis

To identify associations between interpretable nucleus image features and genotype, we conducted an imageQTL analysis. For this analysis, we employed a genome-wide association study approach, using the general linear models implemented in PLINK v2.0 (*48*). For each nucleus feature and GTEx tissue, a set of imageQTLs was calculated. The mode value of a given nucleus feature was used to represent each WSI, treating it as a continuous phenotype across donor samples. Gender, age, death Hardy (which represents the speed of death: 0 = ventilator case, 1 = volent and fast death, 2 = fast death from natural causes, 3 = intermediate death, 4 = slow death), and 20 genotype principal components were included in the imageQTL analysis as covariates to account for population stratification. GTEx genotype data were obtained (file name: ‘GTEx_Analysis_2017-06-05_v8_WholeGenomeSeq_838Indiv_Analysis_Freeze.SHAPEIT2_phased.MAF01.vcf.gz’) and any SNP with a minor allele frequency smaller than 0.01 was excluded from the analysis. The number of individual samples for each tissue type is listed in **SI Appendix, Table S1**. A genome-wide significance threshold of p-value < 5e-8 was applied, and significant SNPs were annotated using Functional Mapping and Annotation of Genome-Wide Association Studies (FUMA) v1.5.2 (**SI Appendix, Dataset S3**) (*49*). We defined overlap between imageQTLs and GTEx eQTLs (**SI Appendix, Dataset S5**) as any imageQTL that is either identical to a GTEx eQTL or in strong linkage disequilibrium with a GTEx eQTL SNP.

### DE Analysis

DE genes were identified in each GTEx tissue and for each type of nucleus feature. The mode value of each nucleus feature was used to represent each WSI. GTEx individuals were categorized into non-overlapping binary groups based on their nucleus features for each tissue and feature type. For example, in thyroid tissue, WSIs were categorized into ‘large’ or ‘small’ nucleus area groups based on the mode of the nucleus area values for each WSI. DE genes were calculated between large and small nucleus area samples in thyroid tissue. This approach was similarly applied to calculate DE genes for other tissues and nucleus features, including nucleus area, circularity, eccentricity, and the nucleus-to-cell area ratio.

We utilized raw bulk gene expression profiles from GTEx (GTEx_Analysis_2017-06-05_v8_RNASeQCv1.1.9_gene_reads.gct.gz) for the DE analysis. Filtering was performed using counts per million (CPM) normalization to remove lowly expressed genes. Genes were excluded if their CPM-normalized expression was > 0.5 in less than 30% of the samples, or if their total raw counts across all individuals were < 500.

DE analysis for each tissue and nucleus feature was conducted using DESeq2’s standard pipeline with the likelihood ratio test (*50*). The analysis included age, gender, genotype ancestry, the death Hardy scale, ischemia time (TRDNISCH), log-transformed unique molecular identifiers as covariates. Contrasts were made between the large and small groups, and multiple testing corrections (Benjamini–Hochberg [BH]) were applied. Genes with an adjusted p-value < 0.01 and abs(log2fold) greater than 0.1 were considered significantly differentially expressed between the contrast conditions. To focus on stronger effects, a more stringent threshold of abs(log2fold) > 0.5 may be applied.

### Epigenome-Wide Association Study for Lung Nucleus Features

Within the GTEx dataset, several tissues have methylation data available, with lung tissue having the largest dataset, consisting of 190 samples across 206,802 CpG sites. The remaining 11 GTEx tissues were not analyzed due to their limited size in methylation data (e.g., muscle; N

= 42). We first divided the samples into binary groups (as described in the DE analysis). EWAS analysis was performed using the ChAMP package (*51*). We calculated the association between methylation values at each CpG site and binary nucleus features. Significant methylation sites were identified using the champ.DMP function, while significant regions were determined using the champ.DMR function. Additionally, we performed GSEA using the internal champ.GSEA function. Multiple hypothesis correction was applied using ChAMP. CpG sites or CpG pathways with an adjusted p-value < 0.05 were considered significant and included.

### HISNUC-X and HISNUC-AGE — Deep Learning Models for Gene Expression and Chronological Age Prediction

#### 1. NIC Image Compression

The NIC framework employs unsupervised learning techniques to compress images into smaller, more manageable forms with minimal information loss. NIC compression can utilize one of three unsupervised learning strategies: autoencoders, contrastive learning, or BiGANs. In our study, we selected BiGANs for autoencoding due to their superior performance.

Gigapixel WSIs ([X, Y, 3]) were divided into adjacent tiles (128×128×3 pixels per tile), and each tile was compressed into a one-dimensional 1D vector (1×1×128) using the NIC algorithm. This process transformed raw pixel data into dense latent space encodings and captured the essential features and patterns while discarding noise and irrelevant information. These 1D vectors were then stacked according to their original spatial arrangement within the WSI, ensuring that spatial relationships between tiles were preserved in the compressed representation. Consequently, the dimensionality of the original RGB image was effectively reduced to [(X/128), (Y/128),128].

#### 2. Preparing Compressed WSIs for Input to HISNUC

Following NIC compression, we obtained compressed WSIs. First, we used a tissue-background segmentation algorithm to effectively remove the background by setting the background pixels to zero. This step was followed by the elimination of entire rows or columns composed solely of empty pixels and focusing the analysis on regions of interest within the WSI. The WSIs were tiled using a sliding window approach, generating 32×32×128 (or 64×64×128) pixel tiles with a 16-pixel (or 32-pixel) step, both column-wise and row-wise. Tiles exceeding a threshold of 450 NaN values (or 1800 NaN) were discarded to maintain data quality. Eight tiles with high tissue occupancy were selected as input. The dataset creation process combined image data, nucleus features, and meta-information, followed by shuffling for randomness, and splitting into training (53%), validation (13%), and testing (33%) sets to rigorously evaluate the model’s performance. We ensured that 33% of the non-training data was used for testing, allowing for the assessment of model generalization.

#### 3. HISNUC Architecture for Gene Expression and Age Prediction

Tiles of compressed WSIs were used as image input for the CNN, followed by fully connected layers. The first convolutional layer had 64 filters and a kernel size of 3×3. These layers were followed by an average pooling layer to reduce the feature map size. A dropout rate of 0.2 was applied thereafter to mitigate overfitting. The architecture included two additional sets of convolutional layers, with 128 and 256 filters, respectively, each followed by average pooling and dropout layers. After the convolutional and pooling layers, the network transitioned to a flattening layer that reshaped the pooled feature maps into a 1D vector. This was followed by multiple dense layers employing ReLU activation. Metadata and nucleus features were processed through dedicated SNNs(*22*). Each encoder consisted of fully connected layers with Scaled Exponential Linear Unit (SELU) activations and AlphaDropout regularization, enabling stable representation learning. The modality-specific embeddings derived from image patches, nucleus features, and metadata information were integrated using a tensor fusion framework, which enables outer product–based interactions and supports effective cross-modal learning (*25*). The resulting fused representation was passed through fully connected layers to perform either gene expression or chronological age prediction, depending on the target task.

For the gene expression prediction model, the output layer comprised 100 units— matching the number of genes being predicted—with a linear activation function to output continuous gene expression levels. In contrast, the chronological age prediction model featured a single-unit output layer with a linear activation function. The model was trained using the SGD optimizer, minimizing mean squared error (MSE), with callbacks for model checkpointing and early stopping to mitigate overfitting. Model evaluation and saving procedures were executed post-training, ensuring the preservation of models with the best validation loss.

We also investigated the potential for multi-instance learning (MIL) (a common WSI embedding technique(*45–47*, *52*, *53*)) to improve WSI encoding with HISNUC. We employed an attention-based MIL architecture to aggregate CNN-compressed features from all available patches per WSI (up to a maximum of 30)(*53*). More specifically, each patch was encoded using the CNN framework described above, and the resulting features were aggregated via an attention-based pooling layer to yield a 128-dimensional slide-level representation. Nucleus features and metadata were encoded separately using the SNN encoders. The modality-specific embeddings and the MIL-aggregated image features were combined via tensor fusion. The resulting fused representation was then passed through fully connected layers to predict the final outcome. The MIL-based HISNUC variant resulted in slightly improved gene expression predictions in a subset of tissues (tibial artery, muscle) compared to the original HISNUC model (**SI Appendix, Fig. S7**).

#### 4. Training and Evaluating the HISNUC-X Model

Tissue-specific genes were calculated using the Tau score (*38*) and defined as those with a Tau score greater than 0.85 and gene expression values exceeding 1. The 100 tissue-specific genes with the highest expression levels were selected. Log-transformed TPM (transcripts per kilobase million) normalized gene expression data from bulk RNA sequencing were utilized as prediction targets.

The gene expression prediction model was trained for each tissue type using varying learning rates. The input tile sizes were typically 32×32×128 for most tissues, though some tissues utilized a tile size of 64×64×128. The nucleus feature inputs also varied by tissue type (**SI Appendix, Dataset S8**). The meta-information input for the HISNUC-X model included height, weight, BMI, age, TRDNISCH, drinking index, smoking index (calculated based on frequency and duration), TRVNTSR (indicating whether the donor was ventilated for less than 24 hours before the estimated procurement start time), gender, race, death Hardy category (DTHHRDY), RNA integrity number (SMRIN), and autolysis score (SMATSSCR). Categorical variables were transformed using one-hot encoding, while numerical variables were normalized. All expanded meta-information was protected data and accessible through dbGAP. For the experiment depicted in **Fig. 4D**, the model was retrained with the “age” variable removed from the meta-information input to eliminate any potential impact on the gene expression prediction results. Training and validation MSE plots were used to track model convergence. Log-scaled loss plots further highlighted subtle changes in training dynamics (**SI Appendix, Fig. S8**). The gene expression level for a sample was estimated by averaging the predictions obtained from all tiles of the WSI for that donor. Evaluation involved comparing predicted and observed gene expression levels from the test set using Pearson correlation coefficients. Three independent experiments were conducted, each with different training, validation, and test sets. The correlation coefficients and p-values were combined using Fisher’s z-transformation and Fisher’s method, with the Holm– Šidák method applied to control for multiple testing.

#### 5. Training and Evaluating the HISNUC-AGE Model

The HISNUC-AGE prediction models were trained by tissue using different learning rates (**SI Appendix, Dataset S10**). The input tile sizes were typically 32×32×128, though some tissues utilized a tile size of 64×64×128. The nucleus feature inputs varied by tissue type. Only nucleus features with age correlations exceeding Pearson’s r = 0.15 were used as input (**SI Appendix, Dataset S10**). The metadata input for the HISNUC-AGE model included height, weight, BMI, TRDNISCH, drinking index, smoking index, TRVNTSR, gender, race, and DTHHRDY. The ‘age’ variable was removed from the meta-information input for all HISNUC-AGE models. The prediction target utilized in the training was precise chronological age rather than categorized into age buckets, which was also protected data and available for download from dbGaP. Training and validation MSE plots were used to track model convergence. Log-scaled loss plots further highlighted subtle changes in training dynamics (**SI Appendix, Fig. S8**).

For chronological age prediction, the donor’s age was estimated by averaging predictions across all tiles from the WSI of the donor. Pearson correlation and c-index were calculated between predictions and targets. MAE was used as an additional metric to assess the performance of the HISNUC-AGE model (**SI Appendix, Dataset S11**). The error bars in **Fig. 6A** represent one standard deviation above and below the mean of the correlation, showing the spread of the data around the average from three independent prediction experiment replicates.

#### 6. SHAP Analysis for Interpreting Feature Importance

We leveraged SHAP to interpret the predictions from our network. SHAP values provided an understanding of image, metadata, and nucleus feature contributions, highlighting which parts of the input are most influential in determining the output. Using GradientExplainer, we generated explanations for a subset of the test images. This method identified the pixels and patterns within the images that significantly influenced the model’s predictions. By plotting these SHAP values, we displayed the impact of each pixel on the model’s output, emphasizing the importance of different regions in the images for predicting the target variable. Additionally, we visualized the contributions of nucleus and metadata features using the SHAP summary and SHAP bar plot functions. This interpretability is crucial for validating the model’s decision-making process and building trust in its predictions. Moreover, it provides potential pathways for further refining our model.

#### 7. Predicting Chronological Age Using Metadata with Elastic Net Baseline Models

To provide a control basis of comparison for our study, we employed elastic net baseline models to make chronological age predictions from datasets consisting of only the metadata with no histological image information, again followed by shuffling for randomness and splitting into training, validation, and testing sets. The meta-information input for the elastic net baseline model included height, weight, BMI, total ischemic time for the donor, drinking index, smoking index, TRVNTSR, gender, race, and DTHHRDY. The training, validation, and testing samples used were identical to those employed in the HISNUC-AGE model, ensuring comparability of the results. Training and testing with several different alpha and L1 ratio values for the elastic net baseline model revealed 0.5 and 0.0226 to be the optimal values, respectively (**SI Appendix, Dataset S12**). The Pearson correlation was computed between the predicted values and the target chronological age (**SI Appendix, Dataset S13**).

#### 8. Comparative Analysis with Multimodal Baselines

A ResNet50 model pretrained on ImageNet was used as an encoder to convert each 224×224 image patch into a 1024-dimensional feature vector(*54*, *55*). Specifically, features were extracted after the third residual block (conv3_block4_out) using spatial average pooling, followed by a dense layer with ReLU activation. These vectors served as high-level representations of morphological patterns within tissue regions. We used an attention-based MIL architecture to aggregate thousands of instance-level patches for each WSI. To integrate different modalities— namely, nucleus features and metadata—with encoded WSI patches, multimodal baselines were constructed using a similar architecture described in the **HISNUC-X and HISNUC-AGE** sections. The overall architecture incorporates SNN encoders for each modality and employs tensor fusion for multimodal integration. These multimodal baselines were applied to the tibial nerve, heart atrial appendages, and testis for both age and gene expression prediction tasks. However, they did not outperform the HISNUC models (**SI Appendix, Fig. S9**).

## Author Contributions

R.M., C.J.F.C, M.B.G. conceptualized the project. R.M., W.Z., C.J.F.C, P.N., X.Z., and M.B.G. contributed to the methodology. R.M., W.Z., C.J.F.C, P.N., X.Z., T.U., and M.B.G. contributed to the investigation. R.M. and M.B.G. contributed to the visualization. M.B.G. contributed to the funding acquisition. R.M., M.B.G. contributed to the project administration. M.B.G. contributed to the supervision. All authors have contributed to the writing and approved of the manuscript.

## Classification

Biological Sciences/Genetics, Computational biology and bioinformatics.

## Acknowledgment

The authors thank Drs. Daniel Palmer and João Pedro de Magalhães (University of Liverpool, UK) for their illustration and help with the tissue-specific gene Tau score calculation. The authors also thank members of the Gerstein Lab, Eric Ni, Declan Clarke, Beatrice Bosari, Matthew Jensen, and Jason Liu, for their valuable discussions and insights during the development of this project and manuscript. The data used for the analyses described in this manuscript were obtained from the GTEx Portal and dbGaP accession number phs000424.v8 on 10/01/2022. This work was supported by the Albert L Williams Professorship funds and the National Institutes of Health (NIH) under Grant U01HG013840.

## Data, Materials, and Software Availability

Code for HISNUC, along with the trained HISNUC-X and HISNUC-AGE models, is available at: https://github.com/gersteinlab/HISNUC under the MIT license. Key resources, including nucleus features extracted with QuPath, imageQTLs, and DE genes, are available through DOI: 10.5061/dryad.8gtht771x.

GTEx-v8 bulk RNA expression and histological data are publicly available through open access at: https://www.gtexportal.org/home/datasets. Genotype data and expanded subject metadata are protected and accessible through dbGAP accession number phs000424.v8.

