## Supplementary Text File for "Machine Learning Models Based on Histological Images from Healthy Donors Identify ImageQTLs and Predict Chronological Age"

**This PDF file includes:**

Figures S1 to S9
Tables S1
Legends for Datasets S1 to S13

**Supplementary Figures**

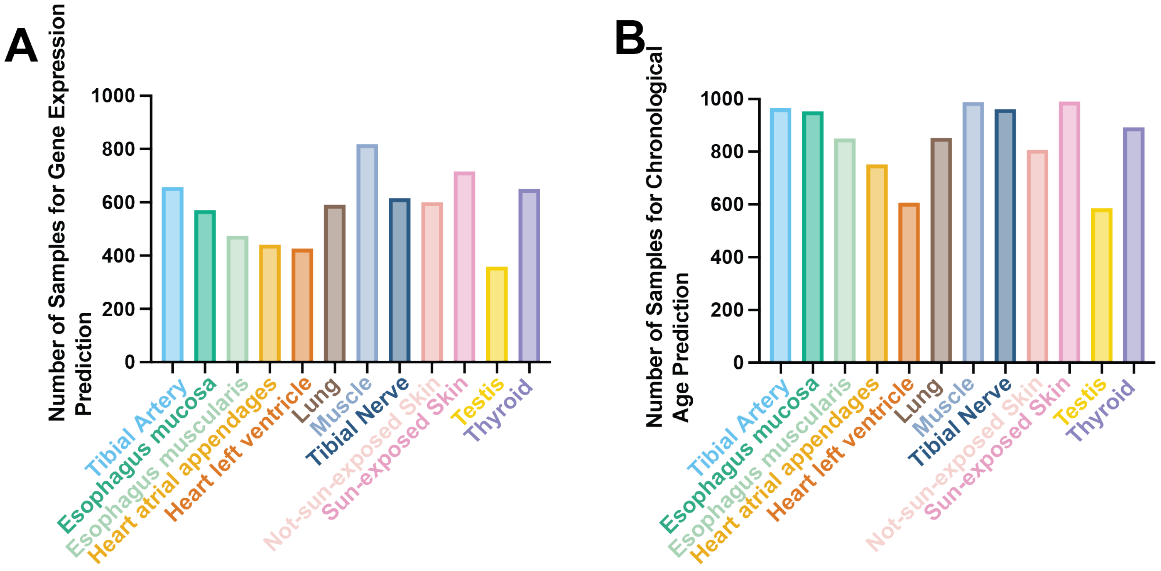


**Fig. S1.** Number of samples used in the chronological age and gene expression prediction models.

Number of samples used for (A) gene expression and (B) chronological age prediction for each GTEx tissue. Muscle, sun-exposed skin, and tibial artery have the highest number of samples, while testis, heart left ventricle, and heart atrial appendages have the lowest number of samples. The number of samples was not indicative of model performance (as shown in **Figs. 4** and **6**).

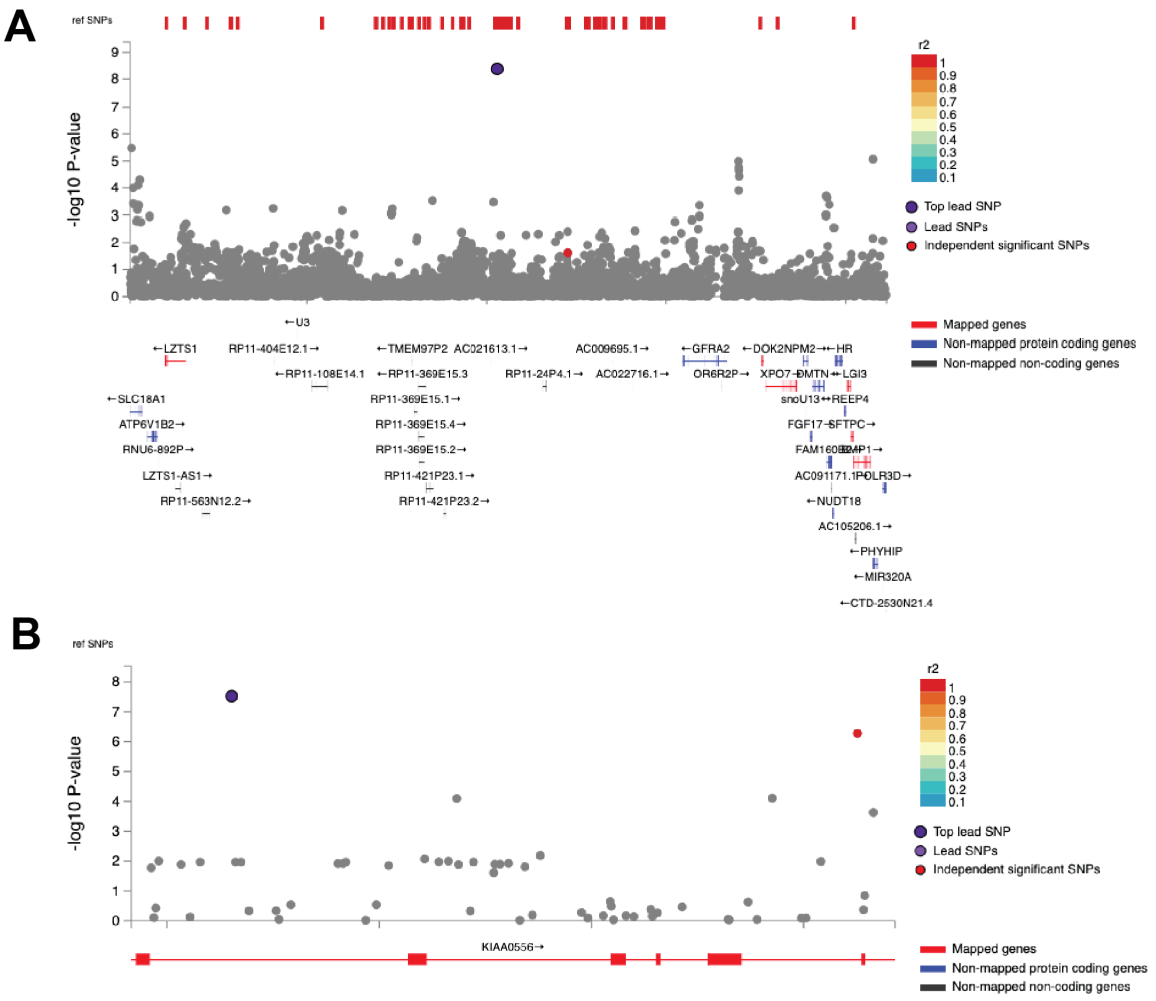


**Fig. S2.** Regional plot for two imageQTL lead SNPs.

Regional plot for top lead SNPs in **Figs. 2C** and **2D** and their mapped genes. Each SNP is color-coded based on the highest $r^2$ to one of the independent significant SNPs. (A) SNP rs76587387 and mapped genes in the GTEx not-sun-exposed skin. (B) SNP rs12444074 and mapped genes in the GTEx tibial nerve.

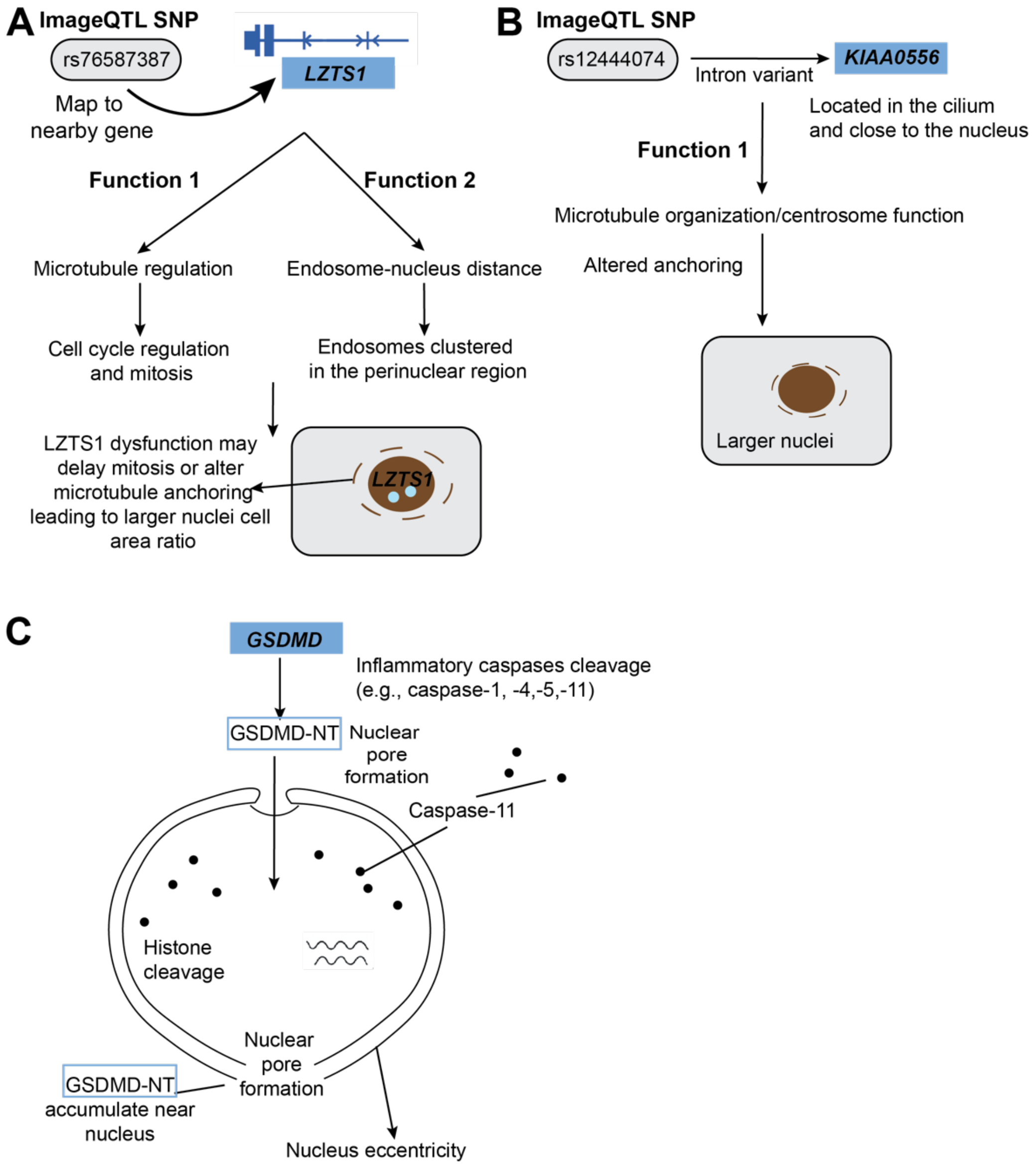


**Fig. S3.** Hypothesized biological mechanisms linking imageQTLs to nuclear morphological features.

(A) *SNP rs76587387* maps near *LZTS1*, which encodes a nuclear-localized tumor suppressor involved in microtubule stabilization, mitotic progression, and regulation of endosome–nucleus positioning—processes that potentially influence nuclear morphology.

(B) *SNP rs12444074* maps near *KIAA0556*, which encodes a microtubule-associated protein localized to the ciliary base and close to the nucleus. Its roles in anchoring microtubules and coordinating nucleus–cytoskeleton interactions suggest potential involvement in regulating nuclear size and positioning.

(C) Schematic illustrating how GSDMD activation disrupts nuclear envelope integrity and alters nuclear morphology through pyroptosis-related membrane pore formation and chromatin remodeling.

A

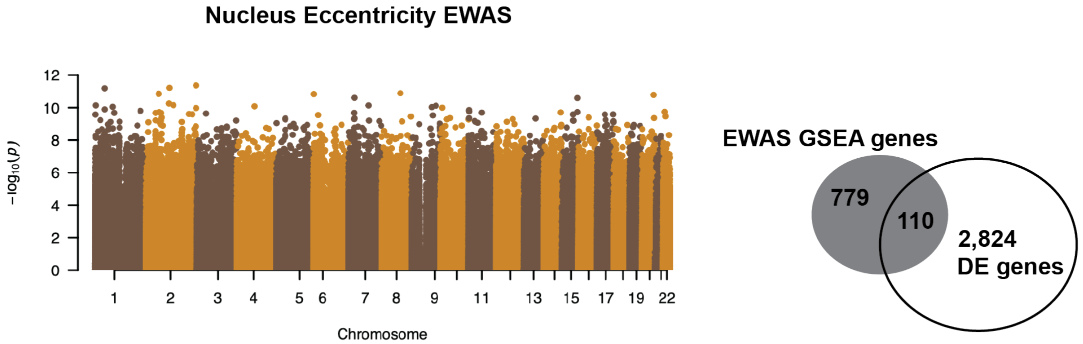


Overlapping genes: 'RGL2', 'MGAT1', 'SPI1', 'SNRPF', 'CHAD', 'CD300A', 'HOXA4', 'FAM65B', 'PADI2', 'ZNF135', 'H2AFY', 'PAPLN', 'ZNF781', 'ST14', 'TWIST1', 'COL11A2', 'GPSM3', 'HOXB3', 'FAM180A', 'RASEF', 'VAX2', 'RADIL', 'C1QTNF7', 'BIN2', 'PDGFD', 'PRRT1', 'GATA5', 'HOXB8', 'RAPGEF3', 'DNASE1L3', 'ABCB1', 'ISM1', 'TNXB', 'GFM1', 'HOXC4', 'NBR2', 'PTK2B', 'ZNF418', 'RPS6KA1', 'HSPB2', 'DOCK2', 'AMPD2', 'TMEM144', 'RAB32', 'CHGB', 'ADAMTS5', 'ADAMTS16', 'DMPK', 'CBFA2T3', 'CYBA', 'DPEP1', 'NFKBIL1', 'WBSCR17', 'NIPAL4', 'LAD1', 'MAGI2', 'HSF4', 'SGIP1', 'MRI1', 'NFATC4', 'ISLR', 'CMTM2', 'ZNF577', 'DIXDC1', 'LST1', 'PIK3R5', 'MSX2', 'MCF2L', 'PRDM6', 'CLDN15', 'ARHGDIB', 'SLC16A3', 'MMP2', 'PPT2', 'FXYD1', 'HOXD9', 'MTMR11', 'CAPG', 'CTSZ', 'SLC6A1', 'KCNAB2', 'HOXA3', 'GATA2', 'EGFL8', 'MYO15B', 'RAP1GAP2', 'ZNF471', 'S100A8', 'GNAI1', 'HOXB2', 'FGR', 'RUSC1', 'FBN2', 'WDFY3-AS2', 'FES', 'SCUBE1', 'JAK3', 'CCRL2', 'ISLR2', 'HLA-DQB1', 'SPC25', 'PCDHA3', 'G0S2', 'TSSK3', 'MOV10L1', 'SLC46A2', 'PGLYRP1', 'TNFRSF10C', 'BASP1','COMP'

B

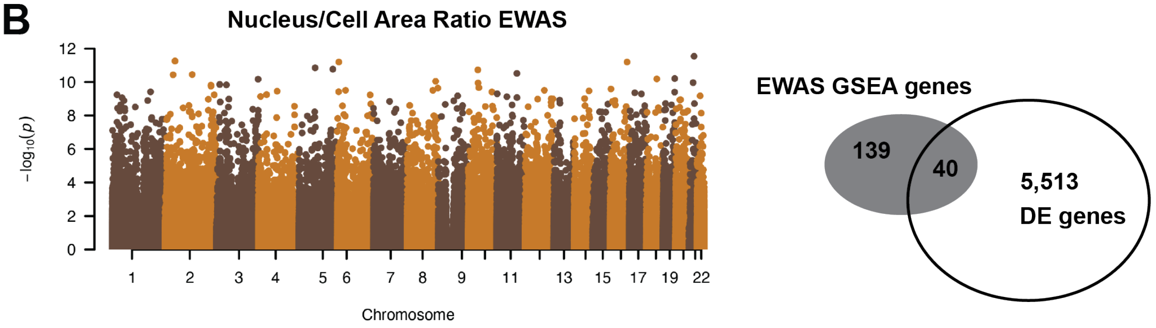


Overlapping genes: 'CTSZ', 'CYBA', 'RUNX1', 'NNAT', 'PCDHGA6', 'PDZRN3', 'ABCC3', 'ZNF471', 'ESRP2', 'KCNIP2', 'C4BPA', 'PAX8', 'ZNF750', 'HTRA4', 'GALNT3', 'MAL2', 'PCDHGA5', 'SSH2', 'GCNT2', 'NRXN3', 'MCCC1', 'HOXD1', 'COL1A2', 'HLX', 'MAP7', 'PGM1', 'PLEKHA2', 'KCNA4', 'CLDN18', 'PCDHGA2', 'PCDHGA4', 'PCDHGA1', 'PCDHGB3', 'ELF5', 'ELF3', 'FBXO39', 'TMEM30B', 'PCDHGA7', 'ATP5G2', 'RNF39'

**Fig. S4.** EWAS Manhattan plots and GSEA analysis for lung.

Manhattan plots for the EWAS between extracted nucleus features and mCpG sites (left). The X-axes of the Manhattan plots represent genomic positions of CpG sites across chromosomes, and the Y-axes represent $-\log_{10}$ P-values. The overlap of the enriched genes from the EWAS GSEA with the DE genes is shown on the right for the GTEx lung tissue dataset (right). Rows represent different features (A) nucleus eccentricity and (B) the nucleus-to-cell area ratio. It should be noted that the EWAS loci identified might not solely reflect differences in nuclear morphology, but could also be influenced by variations in cell type composition. The highlighted overlapping genes may be involved in regulating the phenotype through both methylation changes and gene expression. No significant associations were identified between methylation sites and either nucleus area or nucleus circularity.

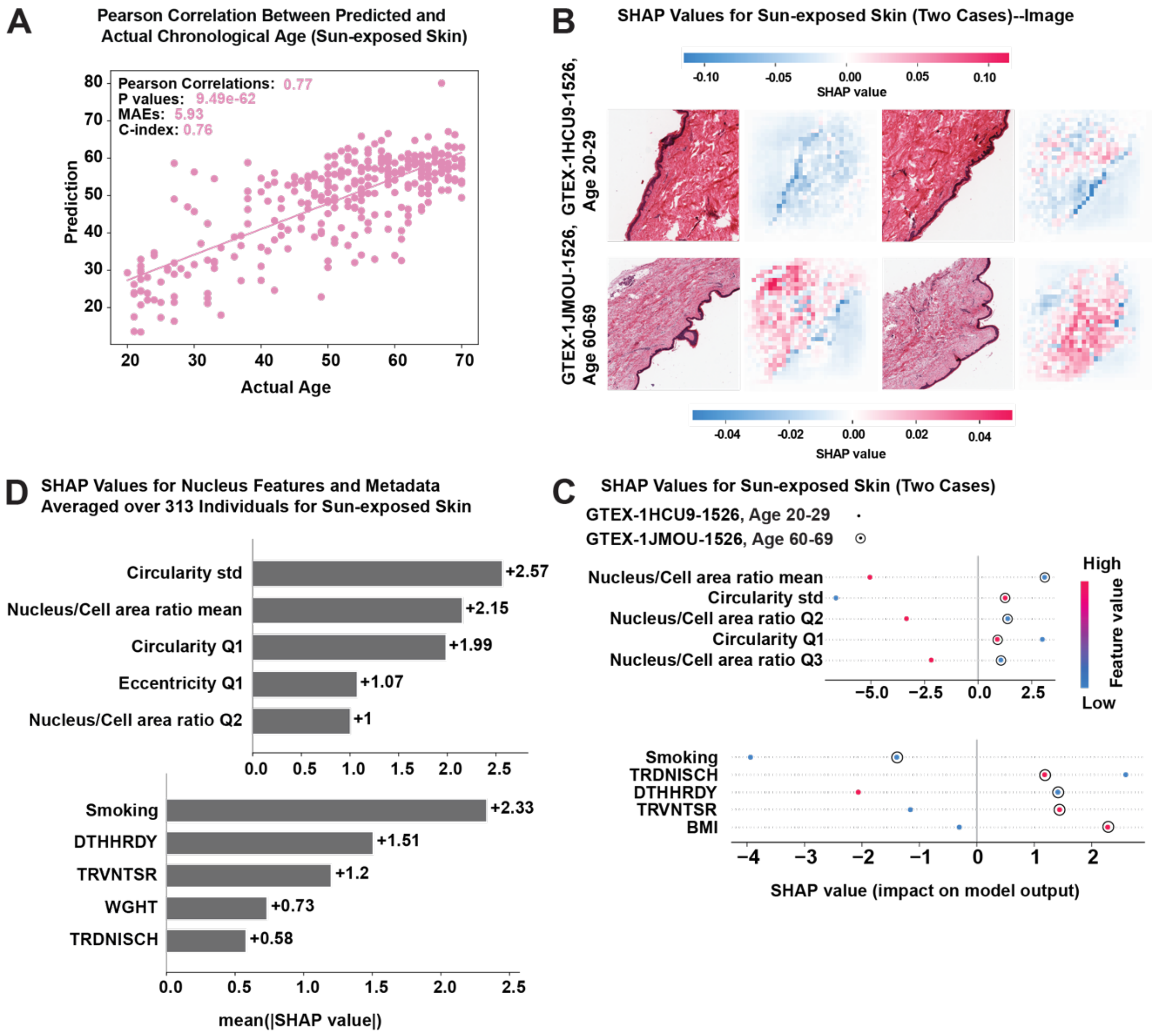


**Fig. S5.** Chronological age prediction for sun-exposed skin using the HISNUC-AGE model.

(A) The scatter plot shows the relationship between predicted age and actual chronological age for sun-exposed skin samples.

(B) SHAP analysis highlights the pixel weights mapped back to the original image for two samples from different age groups, with positive SHAP pixels shown in red (contributing to older age predictions) and negative SHAP pixels shown in blue (contributing to younger age predictions).

(C) Comparison of SHAP values for vectorized features (nucleus features and metadata) between two samples from different age groups. TRVNTSR indicates whether the donor was ventilated for less than 24 hours before the estimated procurement start time, and DTHHRDY represents the speed of death (0 = Ventilator case, 1 = Violent and fast death, 2 = Fast death from natural causes, 3 = Intermediate death, 4 = Slow death). Feature values are colored from high to low, ranging from red to blue. The SHAP values on the X-axis indicate their impact on the model output. Only the top 5 features (ranked by feature importance based on these two samples) that contributed to the model are displayed, distinguishing the two samples from different age groups. TRDNISCH represents total ischemic time for the donor.

(D) Mean absolute SHAP values for vectorized features across the entire test set (N = 313 donors), ranked by feature importance. The smoking index was calculated based on the frequency and duration of smoking.

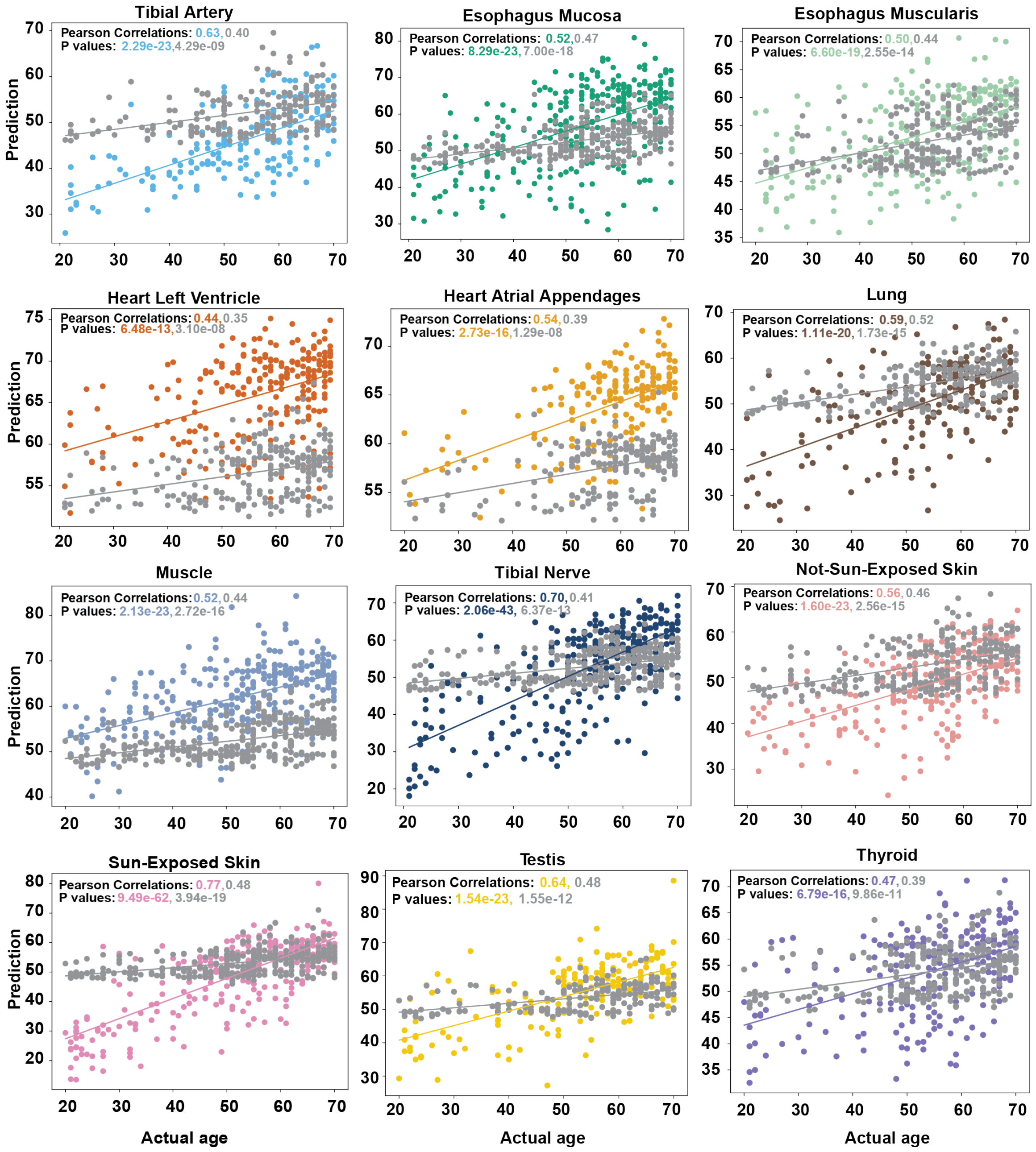


**Fig. S6.** Predicted vs. actual chronological age across tissues.

Scatter plot of the predicted age and the actual chronological age for each tissue from a single prediction experiment with the best-fit line. The scatter plot colored in grey shows the prediction from metadata using the elastic net baseline model as a control. The scatter plot colored by tissue shows the prediction from the HISNUC-AGE models.

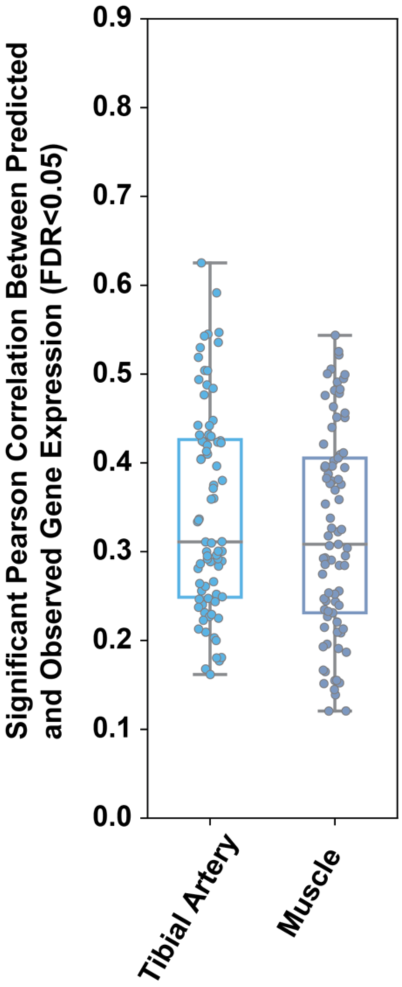


**Fig. S7.** Incorporating MIL slightly improves HISNUC-X gene expression prediction performance in the tibial artery and muscle.

Distribution of Pearson correlations between predicted and actual gene expression levels for the HISNUC-X model with multi-instance learning (MIL) in the tibial artery and muscle.

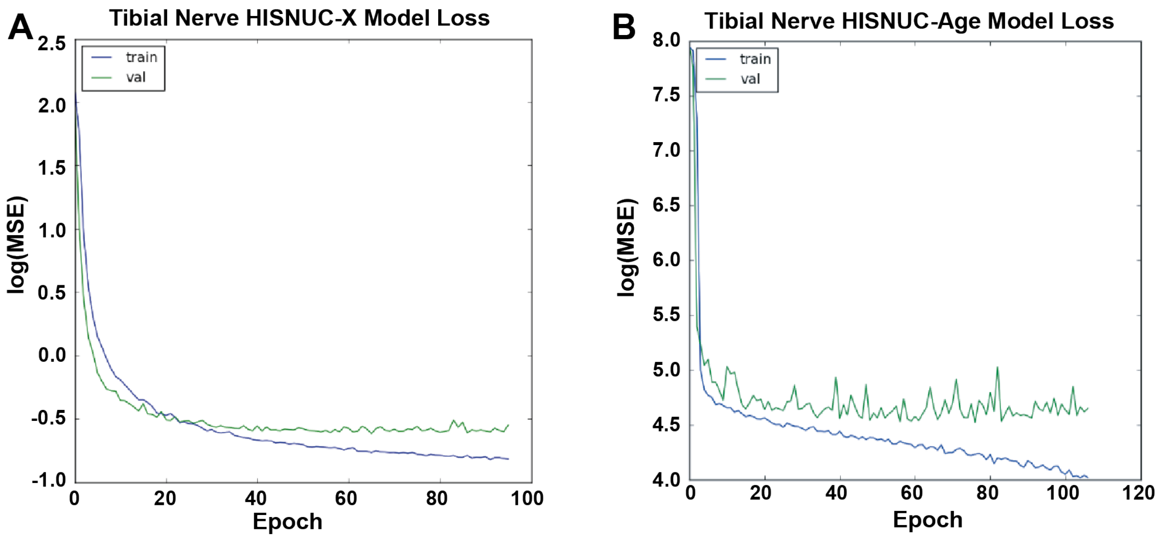


**Fig. S8.** Model loss over training epochs for the HISNUC-X and HISNUC-AGE models on tibial nerve tissue.

(A) Training and validation loss (log(MSE), natural logarithm) across 80 epochs for the HISNUC-X model. (B) Training and validation loss (log(MSE), natural logarithm) across 100 epochs for the HISNUC-AGE model. In both panels, the training curve is colored in blue, and the validation curve is shown in green. The loss decreased significantly at the beginning of training, followed by stabilization. Tibial nerve tissue is presented here as a representative example, with similar trends observed in other tissues.

A

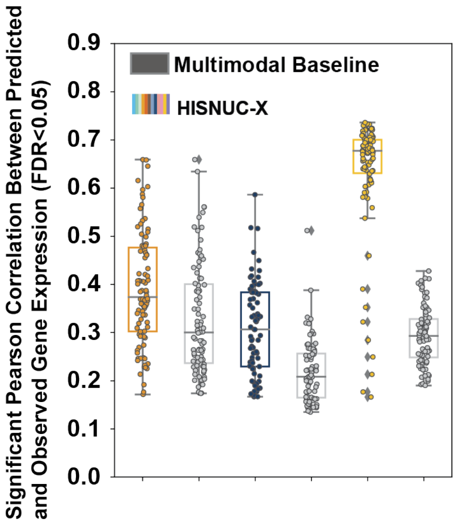


B

| Model | HISNUC-AGE | | | Multimodal Baseline | | |
|---|---|---|---|---|---|---|
| | MAE | Pearson correlation | c-index | MAE | Pearson correlation | c-index |
| Heart atrial appendages | 6.865 | 0.506 | 0.667 | 6.376 | 0.303 | 0.592 |
| Tibial Nerve | 6.933 | 0.694 | 0.736 | 8.842 | 0.331 | 0.621 |
| Testis | 7.117 | 0.618 | 0.713 | 10.125 | 0.327 | 0.604 |

**Fig. S9.** Comparison between HISNUC models and multimodal baselines.

(A) Distribution of Pearson correlations between predicted and actual gene expression levels for the HISNUC-X model and ResNet50-based multimodal baselines. Correlations (adjusted p-value $< 0.05$) were computed across the test set for each gene. Each dot represents a gene, color-coded by tissue type. (B) Performance metrics—including MAE, Pearson correlation, and concordance index (C-index)—for age prediction using the HISNUC-AGE model versus the ResNet50 multimodal baseline.

**Supplementary Tables**

**Table S1.** Data Resource.

| Tissue name | ImageQTL | | HISNUC-X | | | HISNUC-AGE | | |
|---|---|---|---|---|---|---|---|---|
| | No. WSIs used | No. ImageQTL | No. samples used* | Input image size (in pixels) | No. output | No. samples used* | Input image size (in pixels) | No. output |
| Tibial Artery | 828 | 21 | 658 | 32x32x128 | 100 | 965 | 32x32x128 | 1 |
| Esophagus Mucosa | 796 | 2 | 571 | 32x32x128 | 100 | 954 | 32x32x128 | 1 |
| Esophagus Muscularis | 719 | 40 | 475 | 32x32x128 | 100 | 850 | 32x32x128 | 1 |
| Heart Atrial Appendages | 515 | 449 | 427 | 32X32x128 | 100 | 607 | 64X64x128 | 1 |
| Heart Left Ventricle | 628 | 43 | 442 | 32X32x128 | 100 | 752 | 64X64x128 | 1 |
| Lung | 708 | 34 | 591 | 32X32x128 | 100 | 853 | 64X64x128 | 1 |
| Muscle | 829 | 4 | 819 | 64X64x128 | 100 | 989 | 64X64x128 | 1 |
| Tibial Nerve | 823 | 32 | 616 | 32x32x128 | 100 | 962 | 32x32x128 | 1 |
| Not-sun-exposed Skin | 687 | 104 | 601 | 32x32x128 | 100 | 808 | 32x32x128 | 1 |
| Sun-exposed Skin | 831 | 28 | 715 | 32x32x128 | 100 | 990 | 32x32x128 | 1 |
| Testis | 501 | 11 | 359 | 32X32x128 | 100 | 586 | 64X64x128 | 1 |
| Thyroid | 768 | 138 | 650 | 32X32x128 | 100 | 893 | 64X64x128 | 1 |

* The number of samples used for HISNUC corresponds to the model input. Some samples were excluded due to non-qualifying tiles.

**Supplementary Data**

**Dataset S1 (separate file).** Summary statistics of QuPath-extracted nucleus features across samples by tissue. (The raw nucleus features extracted by QuPath are also available as data resources; see Code and Data Availability for details.)

**Dataset S2 (separate file).** Number of ImageQTLs for Each Nucleus Feature and Tissue.

**Dataset S3 (separate file).** ImageQTL Annotated Using FUMA.

**Dataset S4 (separate file).** ImageQTL-Mapped Genes with Potential Roles in Regulating Nuclear Morphology.

**Dataset S5 (separate file).** Overlap between ImageQTLs and GTEx eQTLs.

**Dataset S6 (separate file).** Overlap Between ImageQTL-Mapped Genes and DE Genes.

**Data set S7 (separate file).** Candidate Nuclear Morphology Regulators Supported by ImageQTL and DE Analyses.

**Dataset S8 (separate file).** Parameter Settings and Nucleus Feature Inputs for the HISNUC-X Model.

**Dataset S9 (separate file).** Performance Metrics of the HISNUC-X Model.

**Dataset S10 (separate file).** Parameter Settings and Nucleus Feature Inputs for the HISNUC-AGE Model.

**Dataset S11 (separate file).** Performance Metrics of the HISNUC-AGE Model and Multimodal Baseline.

Column 1 presents the MAE difference between the predicted and actual age. Columns 2–3 show the combined correlation and p-value between the predicted and the actual age across three experiments using the HISNUC-AGE model. Column 4 reports the C-index. Columns 5–8 show the corresponding metrics for the multimodal baseline.

**Dataset S12 (separate file).** Parameter Settings for the Elastic Net Baseline Models.

**Dataset S13 (separate file).** Performance Metrics of the Elastic Net Baseline Models.

Code for HISNUC, along with the trained HISNUC-X and HISNUC-AGE models, is available at: https://github.com/gersteinlab/HISNUC under the MIT license. Key resources, including nucleus features extracted with QuPath, imageQTLs, and DE genes, are available through DOI: 10.5061/dryad.8gtht771x.
